# An anatomical hotspot for striatal dopamine-acetylcholine interactions during reward and movement

**DOI:** 10.64898/2026.01.20.700614

**Authors:** Safa Bouabid, Mai-Anh T. Vu, Christian Noggle, Stefania Vietti-Michelina, Katherine Brimblecombe, Nicola Platt, Liangzhu Zhang, Anil Joshi, Stephanie Cragg, Mark W. Howe

## Abstract

Dopamine (DA) and acetylcholine (ACh) are key neuromodulators that regulate striatal circuits underlying movement and reinforcement learning. Evidence suggests that DA and ACh systems interact, but where and how interactions are expressed across striatal regions in behaviorally relevant release dynamics remains unknown. We applied micro-fiber arrays to simultaneously measure striatum-wide DA and ACh in behaving mice, revealing an anatomical organization in which DA-ACh anti-correlations were concentrated in a hotspot in the anterior dorsolateral striatum (aDLS). Anti-correlations resulted from temporally coincident pairs of spontaneous and event-locked transient peak and dip events occurring in a DA→ACh sequence. The aDLS localized hotspot was consistently expressed within distinct signals associated with unpredicted rewards, learned and extinguished Pavlovian cues, and locomotion initiation and invigoration phases, for which we revealed novel, opposing DA-ACh dynamics. Optogenetic activation of DA neurons selectively suppressed spontaneous ACh release within the aDLS hotspot, and *ex vivo* recordings revealed enhanced D2-mediated ACh inhibition in aDLS relative to ventral regions, suggesting a mechanistic basis for this spatial specificity. These findings demonstrate that DA-ACh interactions during behavior are spatially organized, rather than uniformly conserved, and shape behaviorally relevant dynamics to regulate region-specific functions in learning and movement control.

## Main

Dopamine (DA) release from midbrain terminals and acetylcholine (ACh) release from striatal cholinergic interneurons (CINs) provide critical neuromodulatory inputs that shape motivation, learning, and action. Converging evidence from recording, manipulation, and clinical studies indicates that DA and ACh signals interact to regulate striatal output and behavior across rapid and extended timescales ^1–6^. A large body of *in-vitro* and *in-vivo* evidence supports antagonistic interactions of these modulators, with DA suppressing ACh release from CINs ^7–11^. However, some studies have indicated that these inhibitory interactions may not be ubiquitously expressed, and others support complementary interactions, indicating that DA-ACh relationships may vary flexibly across striatal regions or behavioral contexts ^12–17^. The functional relevance of DA and ACh interactions has primarily focused on two behavioral domains: movement modulation and reward reinforcement learning^18–24^, which engage distinct striatal territories and neuromodulator dynamics. However, prior approaches have not permitted a comprehensive investigation of how DA-ACh signals coordinate across striatal space and time during relevant behaviors, so it remains unresolved whether interactions are uniform or selective to region-specific dynamics.

Antagonistic DA-ACh interactions were first implicated in movement regulation through clinical observations that muscarinic receptor antagonists alleviate motor symptoms in Parkinson’s disease (PD). These findings motivated the influential hypothesis that DA and ACh exert opposing influence on striatal circuits and behavior ^1,25–27^. DA loss in PD is thought to disinhibit CINs, resulting in an anti-kinetic ‘hypercholinergic’ state ^1,21,26–37^. An inhibitory influence of DA on CIN firing and ACh release via CIN D2 receptors has since been demonstrated both *in vivo* and *in vitro* ^3,7–9,11,17,38,39^. Measurements of striatal DA and ACh signals in behaving rodents identified rapid fluctuations aligned to phases of spontaneous locomotion^6,40–42^, which may contribute to movement initiation, vigor, and/or motor learning on rapid or extended timescales^43–46^. However, reports of DA movement signals are highly variable, with some studies describing initiation-linked dips, others peaks, and others no change at all^6,40,45,47–49^. Some of this disagreement likely is due to the fact that DA signals vary across neurons and their downstream striatal regions ^41,43^, and standard approaches typically sample from only 1-2 sites in a given experiment. As a result, how DA and ACh signals co-vary across striatal space during movement, and whether their interactions follow consistent or region-specific patterns, remains unresolved. Studies of DA and ACh interactions in Pavlovian and instrumental conditioning paradigms have also largely indicated antagonistic roles. Striatal CINs exhibit a characteristic multiphasic response to rewards and predictive cues, consisting of a short latency burst followed by a pause in firing (the ACh dip), whereas DA neurons produce transient bursts believed to encode positive reward prediction errors (RPEs). DA lesions or D2 downregulation reduces or abolishes the CIN pause, supporting a model in which DA RPEs inhibit CIN activity via D2 receptors ^3,8^. Both DA peaks and ACh dips are implicated in synaptic plasticity that underlies reinforcement learning, supporting the idea of coordinated, antagonistic control^5,50–52^. Recent work, however, has indicated that under some conditions, CIN D2 receptors only weakly modulate or do not influence the expression of the ACh dip (e.g. to reward ^8,42^), indicating that inhibitory DA-ACh relationships may vary across behavior contexts and/or striatal regions ^9,12,13,15,16,42^. Furthermore, some evidence suggests that positive DA-ACh interactions may dominate in some circumstances ^15,53–56^, perhaps indicating distinct modulatory mechanisms or correlated inputs^9,42,57^. These findings suggest that DA-ACh coupling may vary with behavioral context or striatal region and may involve multiple circuit mechanisms.

Here we address two central gaps: how DA and ACh signals are coordinated across time and striatal space during behavior, and whether their interactions follow uniform principles or exhibit regional or behavior-specific structure. If DA-ACh coupling is consistent across the striatum, it would suggest a common neuromodulatory mechanism acting broadly on striatal computations. Conversely, regional or behavioral specificity would indicate that DA and ACh shape circuit dynamics and behavior through locally tuned interactions. Defining the spatial and temporal organization of DA-ACh coordination is therefore essential for understanding how neuromodulatory systems regulate learning and action and for identifying circuit-level vulnerabilities relevant to disorders that differentially affect distinct neuromodulator pathways.

## Results

### Spatially organized coupling of transient DA-ACh dynamics during behavior

We used large scale multifiber arrays of 46-58 optical micro-fibers (50 µm diameter) distributed across the three-dimensional striatal volume^58^, to simultaneously measure striatum wide DA and ACh dynamics in head-fixed mice (n = 5) spontaneously behaving on a linear treadmill and co-expressing the fluorescent sensors GRAB-rDA3m^59^ and GRAB-ACh3.0^60^ (Fig. 1a,b). Both positive and negative correlations were observed in DA-ACh cross-correlations at single sites and were present at different relative time-lags (Fig. 1c-e). For each site, we defined the ‘dominant’ correlation as the highest magnitude Pearson’s correlation coefficient (the minimum or maximum) calculated within a 1s cross-correlation window (Fig. 1c,d). The polarity of the strongest correlation varied significantly across striatal sites, with some exhibiting dominant positive correlations (42/174 sites, 31 with ACh leading) and others dominant negative (132/174 sites, 127 with DA leading) (Fig. 1e). Dominant negative correlations were most frequent for DA leading time-lags and dominant positive correlations were associated with ACh-leading time-lags (Fig. 1e). To determine whether correlations were anatomically organized across the striatum, we plotted maps of dominant correlation coefficients. These maps revealed a clear topography, with dominant negative correlations concentrated in the anterior dorsolateral striatum (aDLS) and dominant positive correlations in the posterior striatum (Fig. 1f). Plotting positive and negative correlation maps independently indicated that both were aDLS concentrated, despite positive correlations being relatively weaker than negative (Extended Data Fig. 1). To quantitatively define regions with the strongest correlations, we performed a hotspot analysis based on local spatial autocorrelations using the Local Moran’s Index, a Local Indicator of Spatial Association (LISA,^61^). This analysis identified a significant ‘hotspot’ in the aDLS where the dominant correlations were highest, and the polarity of the strongest correlations were negative (Fig. 1f-h). The presence and anatomical localization of this hotspot was remarkably consistent across mice (Extended Data Fig. 2). Mice (n = 3) expressing non-functional red and green fluorescent proteins showed only sparse, weak positive correlations (likely due to correlated broad spectrum noise), and these correlations were not anatomically organized (Extended Data Fig. 1l-k). In sum, these data indicate that relationships between spontaneous DA-ACh dynamics during behavior are not uniform but are temporally specific and spatially concentrated.

**Figure 1:**
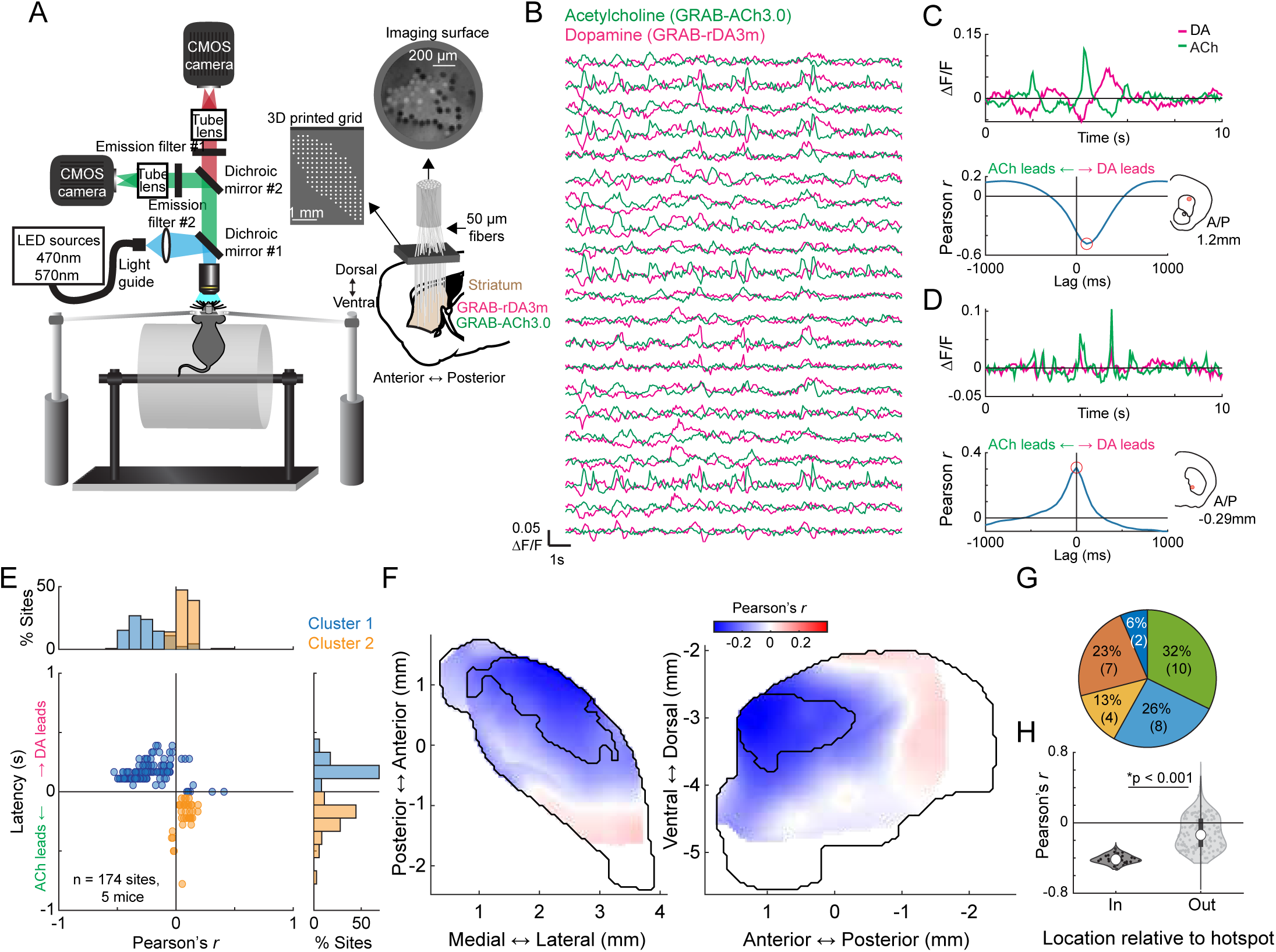
A striatal hotspot in the aDLS for dominant DA-ACh anti-correlations in spontaneous release. **a,** Schematic of the multifiber array approach for striatum-wide measurements of DA and ACh release dynamics in head-fixed, behaving mice. **b,** Example DA and ACh fluorescence traces (ΔF/F) from 21 sites in a single mouse. **c,** Top, example ΔF/F traces from a site exhibiting a dominant negative DA-ACh correlation. Bottom, DA-ACh cross-correlation for the same site across the entire recording. Insets indicate the site location in the coronal plane. **d,** Same as c for another site in the same mouse with a dominant positive DA-ACh cross-correlation. **e,** Dominant DA-ACh cross-correlation coefficient (Pearson’s *r* largest magnitude, i.e., max or min) vs latency for each site (dot) across all mice (5 mice, 174 sites). Clusters identified by k-means. Histograms show the percent of total sites in each cluster binned across correlation coefficients (top) and latencies (right). **f,** Smoothed maps (axial, left; sagittal, right) of dominant DA-ACh cross-correlation coefficients across striatum sites. Black contours delineate a statistically significant aDLS hotspot (Local Moran’s *I*) for dominant correlations. **g,** Contribution (percent) by mouse of total sites in the aDLS hotspot in f. **h,** Violin plot of DA-ACh correlation coefficients for sites inside and outside the aDLS hotspot. Each dot is one site; white circles, median; thick bar, interquartile range; thin lines, 1.5x interquartile range. Sites inside the hotspot show significantly stronger negative correlations (*p=1.67e-16, Wilcoxon rank-sum test).

DA and ACh dynamics often occurred as transient peaks or dips, likely reflecting the coordinated dynamics of cholinergic interneurons or DA terminals (Extended Data Fig. 3a-c, ^6,42^). Therefore, correlation strengths across sites could be determined by the frequency of co-occurring transient events, relationships between the magnitudes of paired events, or both. Correlation strengths varied moment-to-moment across sites and were correlated with the presence of DA and ACh transients, confirming that relationships were not continuous but were dependent on the expression of discrete transient events (Extended Data Fig. 3a-c). We reasoned that co-occurring DA and ACh transients would be most frequent in the aDLS hotspot, where correlation strengths were maximal (Fig. 1f). Four event-pairs categories were considered for transients co-occurring within a 1s window: (1) ACh troughs with DA peaks, (2) ACh peaks with DA troughs, (3) ACh peaks with DA peaks, and (4) ACh troughs with DA troughs (Fig. 2a). Consistent with the correlation results (Fig. 1e), opposite signed pairs (cases 1 and 2) had timings indicating DA leading, while timings of same sign pairs (cases 3 and 4) indicated ACh leading (Fig. 2a). The frequencies of all paired events were highest in the aDLS within a region significantly overlapping with the hotspot identified by correlation analysis, even after correcting for chance pairing rates due to higher aDLS event frequencies (Fig. 2b; Extended Data Fig. 3d-h; p<0.01 for all pairs, volume overlap test, see Methods). Relatively few transients were present with no spatial organization in mice expressing non-functional red and green fluorophores, indicating that the observed effects were not due to hemodynamic or motion artifacts (Extended Data Fig. 3f).

**Figure 2:**
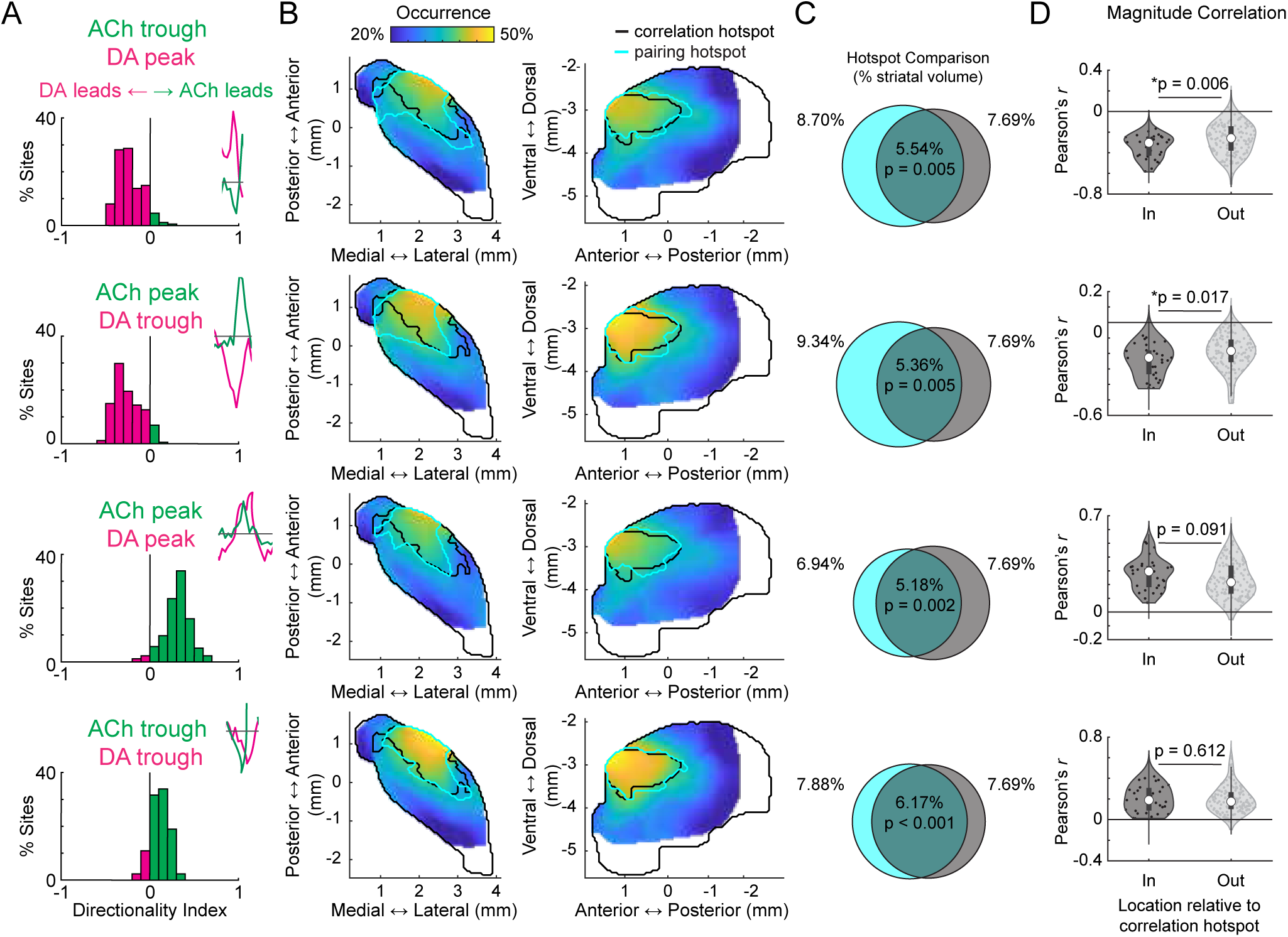
Pairs of coincident DA and ACh transients exhibit polarity dependent temporal relationships and are aDLS concentrated. **a,** Histograms of directionality indices across all sites and mice (5 mice, 174 sites) for each of four spontaneous DA-ACh transient pair types (example ΔF/F traces in inset). Directionality index indicates the extent to which the relative order of events is biased one way or the other, e.g., DA-ACh vs ACh-DA (see Methods). Note the strong bias towards DA leading timings (negative directionality indices) for opposite-polarity pairs and ACh leading for same-polarity pairs. **b,** Smoothed maps (axial, left; sagittal, right) of paired transient frequencies across striatum sites for each transient pair type indicated in the corresponding row of a. Black contours delineate the hotspot identified from global cross-correlation coefficients (Fig. 1f) and cyan contours delineate significant hotspots for the occurrence of each set of transient pairs. **c,** Venn diagrams showing the spatial overlap between the cross-correlation hotspot (black contours in b and Fig. 1f) and the pairing hotspot (cyan contours in b) for each set of transient pairs. Percentages indicate the fraction of the total single-hemisphere striatum volume occupied by each individual hotspot in c (cyan left, black right) and for the intersection of the two hotspots (middle). P-values derived from volume overlap test (Methods). **d,** Violin plots showing correlations between the DA and ACh peak or trough magnitudes across all transient pairs for each site inside or outside the aDLS global correlation hotspot (Fig. 1f). Pair categories match corresponding rows in a. Each dot is one site; white circles, median; thick bar, interquartile range; thin lines, 1.5x interquartile range. P-values, Wilcoxon rank-sum test; *p<0.05. To account for the difference in pair occurrence frequency across sites, the correlation for each site was calculated as the mean coefficient of 10000 repetitions of magnitude correlation for 100 randomly selected transient pairs.

If the DA-ACh correlations were indicative of direct interactions or linked circuit processes (such as common input or feedforward inhibition), the magnitudes of individual event pairs across occurrences would be correlated. For opposite sign event pairs, a majority of sites had significant negative magnitude correlations (e.g. larger peak DA→deeper ACh trough; 121/174 sites significant for ACh troughs vs DA peaks; 81/174 sites for ACh peaks vs DA troughs; Pearson correlation p<0.05; none with significant positive correlation), whereas for same sign pairs, a majority had positive correlations (97/174 significant for peak-peak correlations, 78/174 for trough-trough; Pearson correlation p<0.05; none with significant negative correlation). Magnitude correlations for opposite sign transient pairs, but not same sign pairs, were significantly stronger for sites inside the aDLS global correlation hotspot region (Fig. 1f) than outside (Fig. 2d; p<0.05 for opposite sign pairs, p>0.05 for same sign pairs, Wilcoxon rank-sum test). In sum, inverse polarity DA and ACh events exhibit consistent temporal relationships and are most strongly coupled in both pairing rate and magnitude in the aDLS, measures that jointly explain the anatomical hotspot observed in global DA-ACh anti-correlations. (Fig 1f).

### Conserved spatial structure of distinct anti-correlated dynamics to unpredicted rewards and Pavlovian cues following conditioning and extinction

Anti-correlated DA-ACh dynamics have been widely reported to rewards and reward predictive cues during Pavlovian conditioning and are believed to contribute to associative learning^8,9,52^. We tested whether the aDLS anti-correlation hotspot observed for spontaneous paired events also manifested in cue and reward evoked release. Unpredicted water rewards evoked characteristic sequences of DA and ACh release at many sites: a short latency ACh peak followed by a dip, and a monophasic DA peak (Fig. 3a). To identify the strength and timing of potential ACh interactions with the DA peak, we correlated peak DA ΔF/F magnitudes with ACh ΔF/F at all time lags within +/-500ms of the DA peak across reward deliveries (Fig. 3b). Consistent with results from spontaneous events (Fig. 1e), sites with dominant negative correlations (114/174 sites) indicated the DA peak leading ACh at a latency aligning with the ACh dip (177.9 ± 96.7ms mean latency), and sites with dominant positive correlations indicated the DA peak lagging ACh (58/174 sites) at a latency aligning with the ACh peak (-41.2 ± 218.9ms mean latency; Fig. 3b; Extended Data Fig. 4). The anatomical organization of these correlations indicated a ‘hotspot’ of dominant negative correlations in the aDLS which significantly overlapped with the hotspot identified from spontaneous events (Fig. 3c,e). Correlations were significantly more negative inside than outside the hotspot (Fig. 3d). The distribution of negative correlation strengths was consistent with the dominant correlation spatial pattern in the aDLS, but positive correlations were diffuse and did not overlap with the aDLS hotspot (Extended Data Fig. 5a,b).

**Figure 3:**
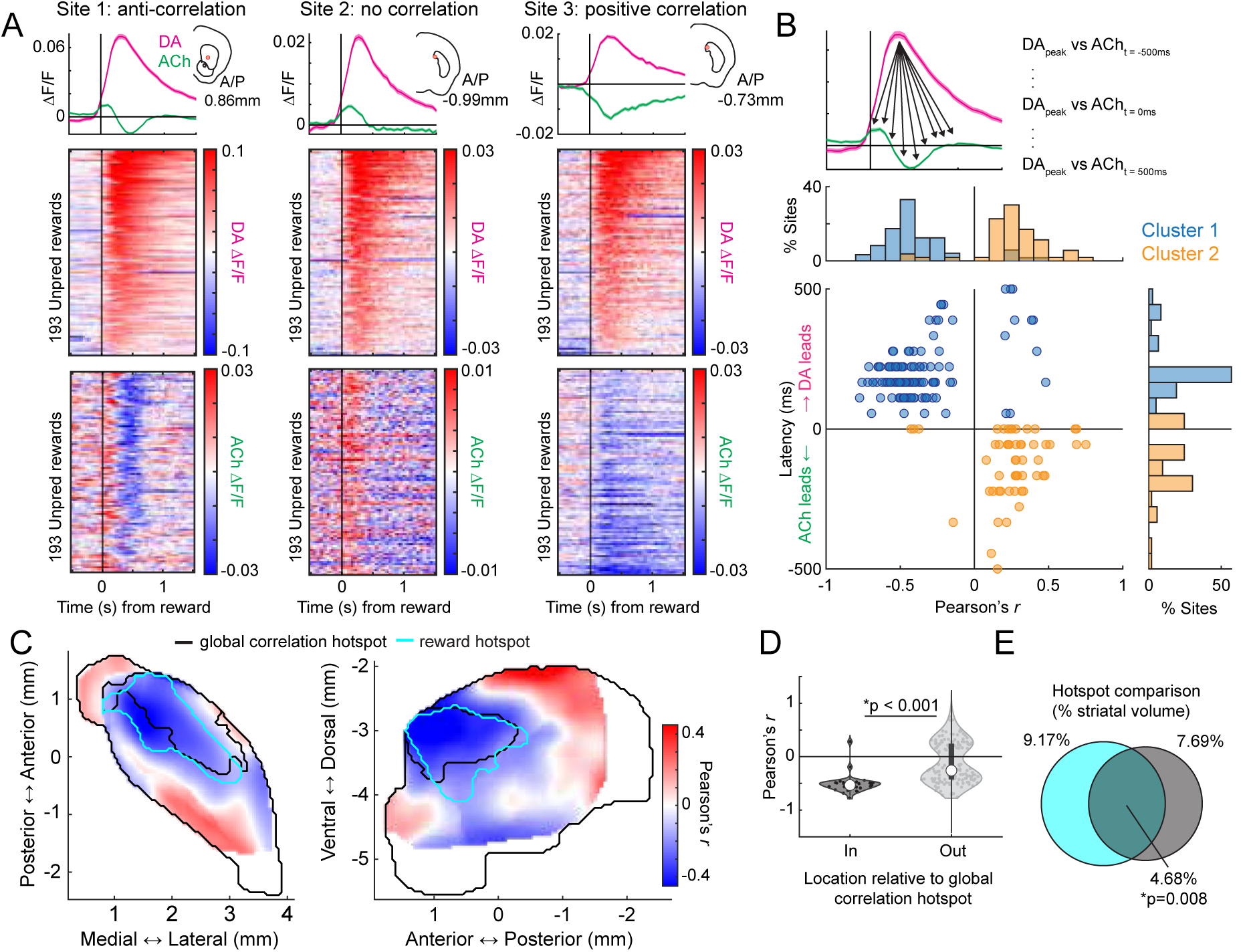
An aDLS localized hotspot for DA-ACh magnitude anti-correlations in unpredicted reward dynamics. **a,** Mean (top) and trial-by-trial (bottom) DA and ACh ΔF/F aligned to unpredicted reward consumption for three example sites in the same mouse with different correlations between the trial-by-trial DA ΔF/F peak magnitudes and ACh (see b, top). Note that despite the presence of prominent dips in ACh ΔF/F for sites 1 and 3, cross-trial magnitude correlations with the DA peak were opposite. Insets indicate the site location in the coronal plane. Shaded regions, S.E.M. **b,** Top: magnitude cross-correlation approach. DA peak ΔF/F magnitudes on each trial were correlated with ACh ΔF/F magnitudes at lags within a +/-500ms window relative to DA peak. Bottom: dominant DA-ACh correlation coefficient magnitude vs latency for each site (dot) across all mice (5 mice, 174 sites). Clusters identified by k-means. Histograms show the percent of total sites in each cluster binned across correlation coefficients (top) and latencies (right). **c,** Smoothed maps (axial, left; sagittal, right) of dominant correlation coefficients for trial-by-trial DA ΔF/F peak vs ACh magnitude (b top) across striatum sites. Black contours delineate the global cross-correlation hotspot (Fig. 1f) and cyan contours delineate significant hotspots for reward anti-correlations. **d,** Violin plots showing dominant DA-ACh reward correlation coefficients (see b) for each site inside or outside the aDLS global correlation hotspot (Fig. 1f). Each dot is one site; white circles, median; thick bar, interquartile range; thin lines, 1.5x interquartile range. P-values, Wilcoxon rank-sum test; *p<0.05. **e,** Venn diagrams showing the spatial overlap between the cross-correlation hotspot (black contour, c and Fig. 1f) and the reward correlation hotspot (cyan contours in c). Percentages indicate the fraction of the total single-hemisphere striatum volume occupied by individual hotspots (cyan left, black right) and for the intersection of the two hotspots (middle, p=0.008, volume overlap test, see Methods).

We next performed the same analyses on cue-evoked DA and ACh dynamics following Pavlovian conditioning and extinction (Fig. 4). Like unpredicted rewards, conditioned cues evoked a temporal sequence of ACh peak→ DA peak→ ACh dip at a majority of sites (49/75 sites). Correlations of the ACh ΔF/F with the peak DA ΔF/F were negative at lagging timepoints (39/75 sites; ACh lagging DA; 202.28 ± 128.4ms) and positive at leads (26/75 sites; ACh leading DA; - 79.1 ± 224.0ms), consistent with the unpredicted reward findings (Fig. 4c). The spatial structure of DA-ACh correlations was also similar, with a significant aDLS hotspot of dominant anti-correlations overlapping with the hotspot for paired spontaneous events (Fig. 4b-e). Positive correlation magnitudes were diffuse and did not exhibit a significant spatial structure (Extended Data Fig. 5c,d).

**Figure 4:**
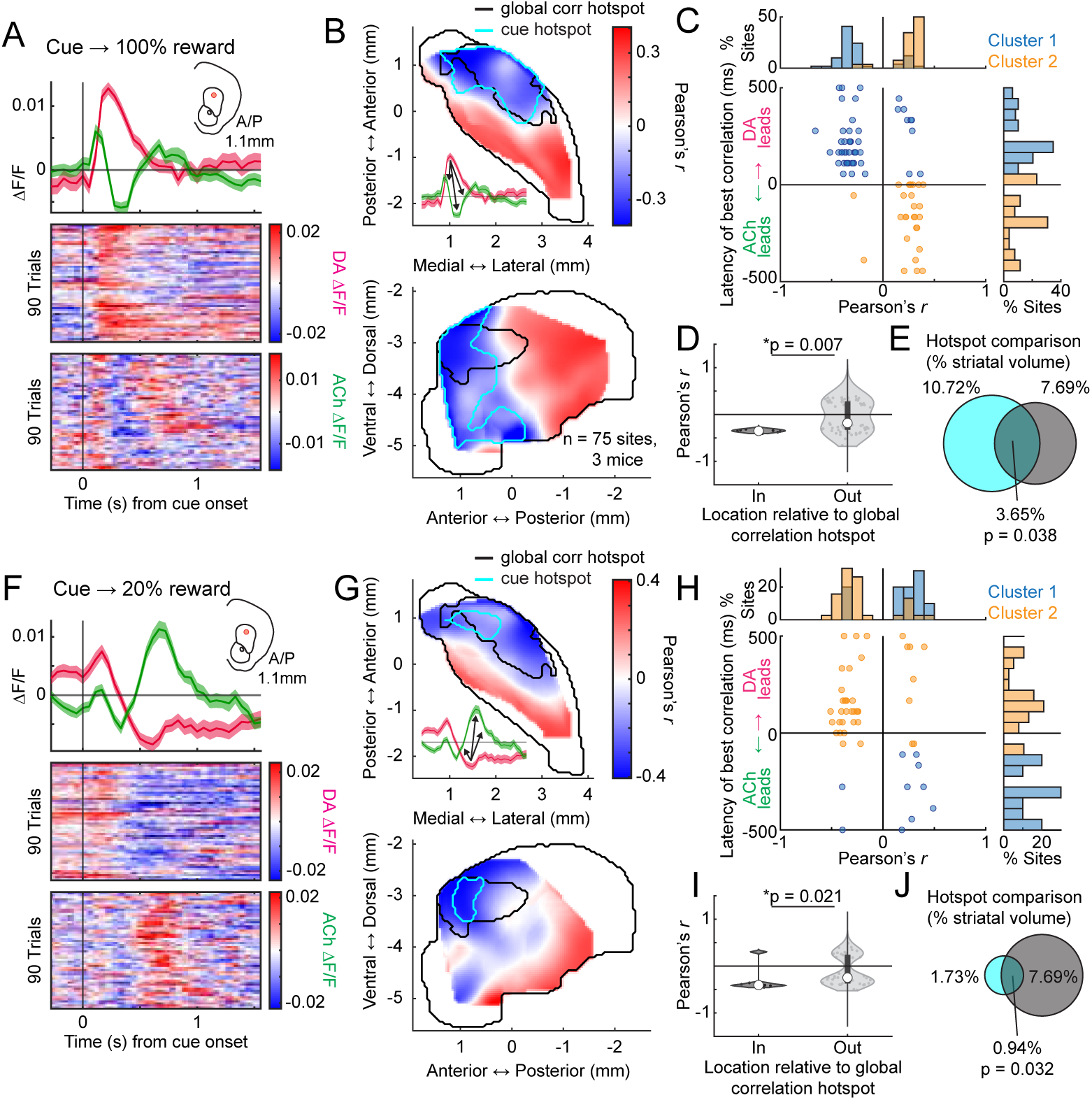
The DA-ACh anti-correlation hotspot in aDLS is consistent for dynamics around conditioned cues and for inverse dynamics following extinction. **a,** Mean (top) and trial-by-trial (bottom) DA and ACh ΔF/F aligned to the presentation of a conditioned light cue associated with upcoming reward (100% probability) after a 3s delay. Inset indicates the site location in the coronal plane. Shaded regions, S.E.M. **b,** Smoothed maps (axial, top; sagittal, bottom) of dominant cross-correlation coefficients (max or min) for trial-by-trial cue-evoked DA ΔF/F peak vs ACh magnitudes at time lags within +/-500ms (cross-correlation schematic inset and Fig. 3b, top) across striatum sites (75 sites, 3 mice). Black contours delineate the global cross-correlation hotspots (Fig. 1f) and cyan contours delineate significant hotspots for conditioned cue anti-correlations. **c,** Dominant conditioned cue-evoked DA-ACh correlation coefficient (see b) vs latency for each site (dot) across all mice. Clusters identified by k-means. Histograms show the percent of total sites in each cluster binned across correlation coefficients (top) and latencies (right). **d,** Violin plots showing dominant DA-ACh conditioned cue correlation coefficients for each site inside or outside the aDLS global correlation hotspot (Fig. 1f). Each dot is one site; white circles, median; thick bar, interquartile range; thin lines, 1.5x interquartile range. P-values, Wilcoxon rank-sum test; *p<0.05. **e,** Venn diagrams showing the spatial overlap between the global cross-correlation hotspot (black contour, b and Fig. 1f) and the conditioned cue correlation hotspot (cyan contours in b). Percentages indicate the fraction of the total single-hemisphere striatum volume occupied by individual hotspots (cyan left, black right) and for the intersection of the two hotspots (middle, volume overlap test, see Methods). **f,** Same as a but for cue-evoked signals on trials following a cue-specific partial extinction where the reward probability was down-shifted to 20%. **g,** Same as b for extinction sessions. Dominant correlation coefficients were obtained via trial-by-trial correlation of DA trough ΔF/F magnitudes with ACh ΔF/F at time lags within +/-500ms of DA trough (cross-correlation schematic inset) for sites with significant DA dip (48 sites, 5 mice). **h-j,** Same as c-d for extinction sessions.

After learning, mice underwent cue-specific extinction training, in which the reward probability associated with one of two conditioned cues was downshifted from 100% to 20% probability of reward delivery (Fig. 4f-h). This paradigm has been previously shown to elicit cue-evoked DA dips and long-latency (‘late’) ACh peaks in the anterior dorsal region, dynamics consistent with negative prediction error encoding (Fig. 4a)^62^. With our simultaneous DA-ACh measurements, we could determine whether the timing and spatial organization of DA dip→ACh late peak dynamics aligned with the inverse DA peak→ACh dip relationship during initial conditioning (Fig. 4a-e). Indeed, the timing relationships for correlations with DA dip magnitudes closely resembled those for the DA peak sites with dominant negative correlations indicating ACh lagging (DA dip→Ach late peak) (Fig. 4h; 41/74 sites; mean lag 117.9 ± 176.2ms). The spatial organization of dominant negative correlations was also aligned, significantly overlapping with the aDLS correlation hotspot (Fig. 4g,i,j; Extended Data Fig. 5e,f). Positive correlations, associated with ACh leading the DA dip, were not significantly spatially concentrated (Extended Data Fig. 5e).

Together, these results indicate a common temporal relationship (DA leading ACh) and anatomical hotspot for DA-ACh anti-correlations occurring spontaneously and associated with behaviorally relevant dynamics to cues and rewards during Pavlovian conditioning and extinction. Positive correlations were also observed but had an opposite temporal relationship (DA following ACh) and did not exhibit a consistent spatial organization across conditions (Extended Data Fig. 8).

### Spatially organized anti-correlations corresponding to phase specific locomotion signals

Imbalances between striatal DA and ACh have been implicated in movement symptoms of Parkinson’s Disease, perhaps through a hypercholinergic state resulting from a loss of ACh suppression by DA^29–37^. Coordinated DA and ACh dynamics have been reported in the dorsal striatum during locomotion, with correlated and anti-correlated signals corresponding to distinct phases of locomotor bouts^6,42^. Despite these findings, it remains unknown how striatum-wide DA and ACh dynamics are organized during locomotion and whether DA-ACh coupling of locomotion related signals is spatially concentrated, as we found for spontaneous and cue/reward related events. To address this, we analyzed simultaneous DA and ACh signals around locomotor initiations. Initiations were carefully selected to ensure little or no prior movement^6,42^ (Methods). Consistent with other work^6,42^, we found that initiations from rest were associated with rapid ACh increases at many sites, followed by decreases (131/174 sites, Fig. 5a-c). In contrast, DA release at some sites (119/174 sites) rapidly decreased, and at others in the same mouse, rapidly increased (73/174 sites) (Fig. 5a-c). Unlike ACh release, only a minority of sites had both DA increases and decreases (32/174 sites). These DA and ACh dynamics occurred at distinct latencies relative to initiation: DA dips were earliest (median latency 111.11, IQR 55.56-708.33 ms), followed by ACh peaks and DA peaks (median latencies 222.22, IQR 222.22-277.78 ms; and 227.78, IQR -388.89-444.44 ms, respectively), then ACh dips (median latency 611.11, IQR 500.00-722.22 ms). This sequential expression of distinct DA and ACh dynamics was topographically organized: early DA dips were concentrated in the aDLS, followed by DA peaks in the dorsomedial and posterior striatum (Fig. 5e,f; Extended Data Fig. 6a). The ACh peaks, though widespread, were strongest in the aDLS, and later ACh dips were localized to the aDLS (Fig. 5e,g).

**Figure 5:**
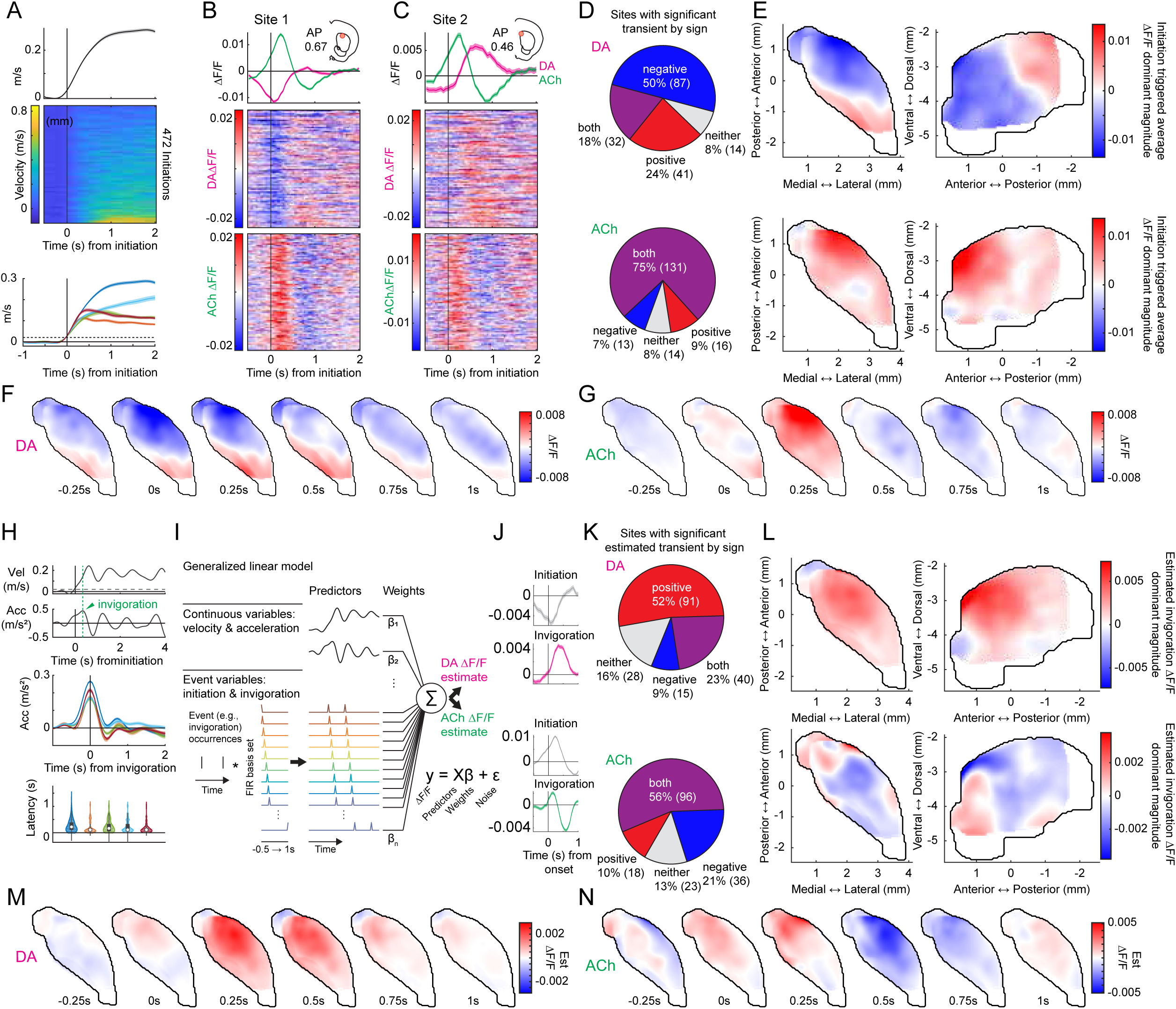
Spatial and temporal organization of DA and ACh signals associated with locomotion initiation and invigoration. **a,** Top: mean (top) and trial-by-trial (raster, bottom) velocity of a single mouse aligned to locomotion initiations, sorted by mean velocity post-initiation. Bottom: mean velocity of all mice (each color represents a single mouse, n = 5) aligned to initiation. Shaded regions, S.E.M. **b,** Mean (top) and trial-by-trial (rasters, bottom; velocity-sorted as in a) DA and ACh ΔF/F aligned to locomotion initiation for an example site exhibiting a significant DA dip. Shaded regions, S.E.M **c,** Same as b for another example site from the same mouse exhibiting a significant DA peak. **d,** Proportion of sites with a significant locomotion initiation-evoked ΔF/F (DA, top; ACh, bottom) by sign (positive, negative, both, or neither) **e,** Smoothed maps (axial, left; sagittal, right) for DA (top) and ACh (bottom) of the dominant ΔF/F magnitude -0.5s to 1s around locomotion initiations. **f,** Smoothed axial maps of DA ΔF/F at 0.25s intervals from -0.25 to 1s from locomotion initiations. **g**, Same as f for ACh. **h**, Top: invigoration events (green dashed line) were defined as the time of local max acceleration immediately following locomotion initiations. Middle: mean acceleration of all mice (each color represents a single mouse, n = 5) aligned to invigoration events. Shaded regions, S.E.M. Bottom: violin plots for each mouse of the latency from locomotion initiation to invigoration event. Each dot is one event; white circles, median; thick bar, interquartile range; thin lines, 1.5x interquartile range. **i,** schematic of the generalized linear model used to estimate the relationship between locomotion variables and DA and ACh ΔF/F (Methods). **j,** GLM-estimated locomotion initiation (gray) and invigoration (magenta or green) evoked ΔF/F (DA, top; ACh, bottom) from the same fiber as in b. **k,** Proportion of sites with a significant estimated invigoration-evoked ΔF/F (DA, top; ACh, bottom) by sign (positive, negative, both, or neither). **l,** Smoothed maps (axial, left; sagittal, right) for DA (top) and ACh (bottom) of the dominant estimated ΔF/F magnitude -0.5s to 1s around locomotion invigorations. **m,** Smoothed axial maps of DA ΔF/F at 0.25s intervals from -0.25 to 1s from locomotion invigorations. **n,** Same as m, but for ACh.

We reasoned that the multi-phasic dynamics observed for locomotion initiations might represent two distinct phases: the initiation from rest and the invigoration into high velocity running (Fig. 5h). Indeed, aligning ΔF/F to the peak acceleration following locomotion initiations on bouts where animals invigorated high velocity locomotion emphasized the DA peaks and ACh dips, whereas aligning to the initiation from rest emphasized the DA dips and ACh peaks (Extended Data Fig. 6b-d). To isolate the ΔF/F associated with invigorations, we modeled the DA and ACh activity around locomotion initiations as a function of movement initiation events, invigoration events, and continuous velocity and acceleration (Fig. 5i). This approach captured DA peaks and ACh dips associated with invigorations, distinct from the DA dips and ACh peaks at initiations (Fig. 5j). DA peaks were topographically organized across the anterior dorsal striatum and preceded the ACh dips which occurred across the same regions (Fig. 5l-n; Extended Data Fig. 6f). In summary, these results reveal a novel, striatum-wide spatiotemporal topography of bi-directional DA and ACh release dynamics associated with locomotion initiations and invigorations.

To investigate direct correlations between initiation and invigoration related DA and ACh dynamics, we took a similar approach to the cue and reward analyses (Fig. 3b). Since the initiation related DA dip occurred earliest, we asked whether DA dip magnitudes correlated with ACh ΔF/F at all other time lags across all initiations for each site. Similar to the cue-evoked DA dips during extinction (Fig. 4h), dominant negative correlations were present between DA and ACh with timings indicating DA leading (Fig. 6a; mean latency 193.80 +/- 177.68ms). As expected, these correlations were concentrated in the aDLS, the area expressing the largest initiation evoked DA dips, and this region significantly overlapped with the previously identified hotspot (Fig. 6b-d). To determine whether the coupling hotspot also held for coupling of inverse DA-ACh signals at invigorations, we correlated the GLM estimated DA peak magnitudes across all invigoration events with ACh at each site. Again, dominant negative correlations were observed at lags relative to the DA peak and were selectively concentrated in the aDLS, significantly overlapping with the previously identified anatomical hotspot (Fig. 6e-h;). Unlike negative correlations, positive DA-ACh correlations were diffuse and not aDLS concentrated (Extended Data Fig. 7).

**Figure 6:**
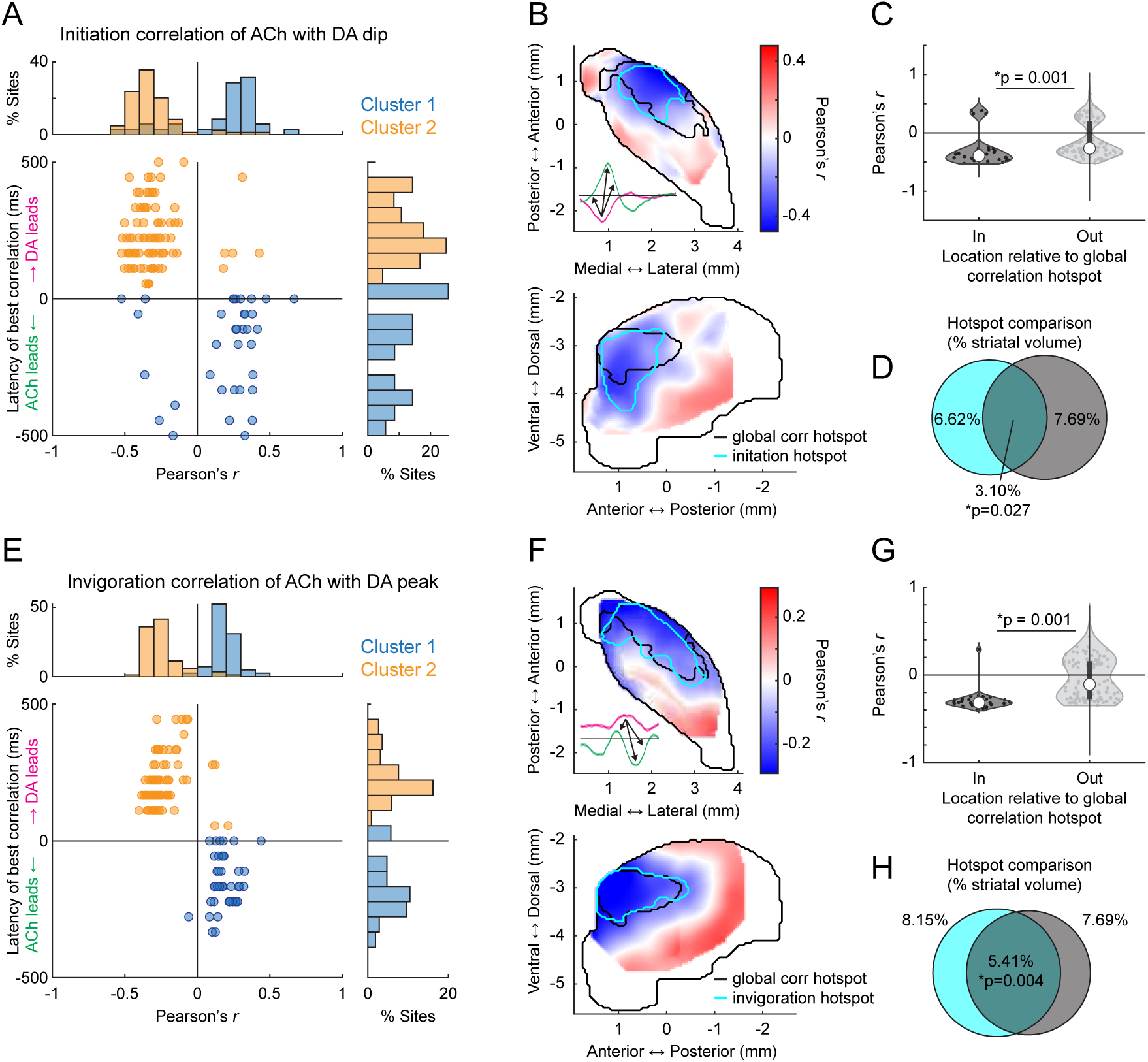
A conserved aDLS hotspot localized for DA-ACh anti-correlations in locomotion initiation and invigoration dynamics. **a,** Dominant locomotion initiation-evoked DA-ACh correlation coefficient (Pearson’s *r* min or max) vs latency for each site (dot) across all mice. Correlations were calculated for trial-by-trial initiation-evoked DA ΔF/F troughs vs ACh magnitudes at time lags within +/-500ms of DA trough (cross-correlation schematic inset in b) across striatum sites with significant DA dips (119 sites, 5 mice). Clusters identified by k-means. Histograms show the percent of total sites in each cluster binned across correlation coefficients (top) and latencies (right). **b,** Smoothed maps (axial, top; sagittal, bottom) of dominant correlation coefficients for trial-by-trial DA ΔF/F peak vs ACh magnitude (b top) across striatum sites. Black contours delineate the global cross-correlation hotspot (Fig. 1f) and cyan contours delineate significant hotspots for initiation anti-correlations. **c,** Violin plots showing dominant DA-ACh locomotion initiation correlation coefficients (cross-correlation schematic inset in b) for each site inside or outside the aDLS global correlation hotspot (Fig. 1f). Each dot is one site; white circles, median; thick bar, interquartile range; thin lines, 1.5x interquartile range. P-values, Wilcoxon rank-sum test; *p<0.05. **d,** Venn diagrams showing the spatial overlap between the cross-correlation hotspot (black contour, b and Fig. 1f) and the initiation correlation hotspot (cyan contours in c). Percentages indicate the fraction of the total single-hemisphere striatum volume occupied by individual hotspots (cyan left, black right) and for the intersection of the two hotspots (middle, P-values, volume overlap test, see Methods). **e-h**, Same as a-d, but for DA-ACh correlations calculated relative to GLM derived invigoration-evoked DA peak magnitudes (131 sites, 5 mice; cross-correlation schematic inset in f).

In summary, our results indicate that movement initiations and locomotion invigorations evoke dissociable patterns of sequential DA and ACh release across distinct striatal domains. DA-ACh anti-correlations associate with leading DA events (DA dip→ACh peak for initiations, and DA peak→ACh dip for invigorations), and negative correlation strength is concentrated in an aDLS hotspot, conserved across paired spontaneous events, Pavlovian conditioning, and movement (Extended Data Fig. 8a-d). Positive correlations, in contrast, were associated consistently with opposite timings (ACh leading) and did not exhibit a robust or consistent spatial topography across behavioral contexts (Extended Data Fig. 8e,f).

### Optogenetic dopamine stimulation drives spatially selective ACh suppression in the hotspot

The consistent aDLS spatial topography and temporal order of DA-ACh anti-correlations across behavioral contexts suggests a possible direct intrinsic origin in which inhibitory modulation of ACh release by DA is preferentially concentrated in the aDLS^14^. Alternatively or in addition, the anti-correlation structure may reflect an indirect circuit interaction, such as feedforward GABAergic inhibition communicated through common glutamatergic inputs. To test whether DA inputs exert heterogeneous modulation of ACh release across the striatum, we optogenetically stimulated midbrain DA neuron cell bodies (SNc/VTA) with ChR2 in DAT-cre mice while simultaneously measuring striatum-wide DA and ACh release with fiber arrays (Fig. 7a, Extended Data Fig. 10a). We confirmed that an aDLS DA-ACh anti-correlation hotspot was present in this separate cohort of mice which significantly overlapped with that reported in previous cohorts (Extended Data Fig. 9a-d). Stimulation light power was calibrated to evoke a change in DA ΔF/F approximately equivalent to average ΔF/F magnitudes measured in the same mice to unpredicted reward deliveries (Fig. 7b,c). Significant stimulation evoked release was measured at all sites but one (99%, 179 of 180 fibers; Extended Data Fig. 10b). In contrast, significant ACh suppression following DA stimulation was observed only in a subset of fibers (64%, 116 of 180 sites), and these responses were spatially concentrated in the aDLS (Fig. 7b-e). The spatial concentration of ACh stimulation-evoked dips significantly overlapped with the anti-correlation hotspot for spontaneous release events, and although suppression was observed at some sites outside the hotspot, ACh ΔF/F dip magnitudes were significantly higher, on average, inside than out (Fig. 7f,g). These findings could not be attributed to spatial heterogeneities in the magnitudes of stimulation-evoked DA release (Extended Data Fig. 10b-e). Therefore, DA neuron stimulation is sufficient to drive ACh dips, independently of behaviorally evoked release, preferentially within the same aDLS hotspot containing the strongest anti-correlations during behavior. Collectively, these results support an intrinsic physiological mechanism for a spatially organized inhibitory modulation of ACh release through the midbrain DA system, which regulates specific behaviorally relevant dynamics.

**Figure 7:**
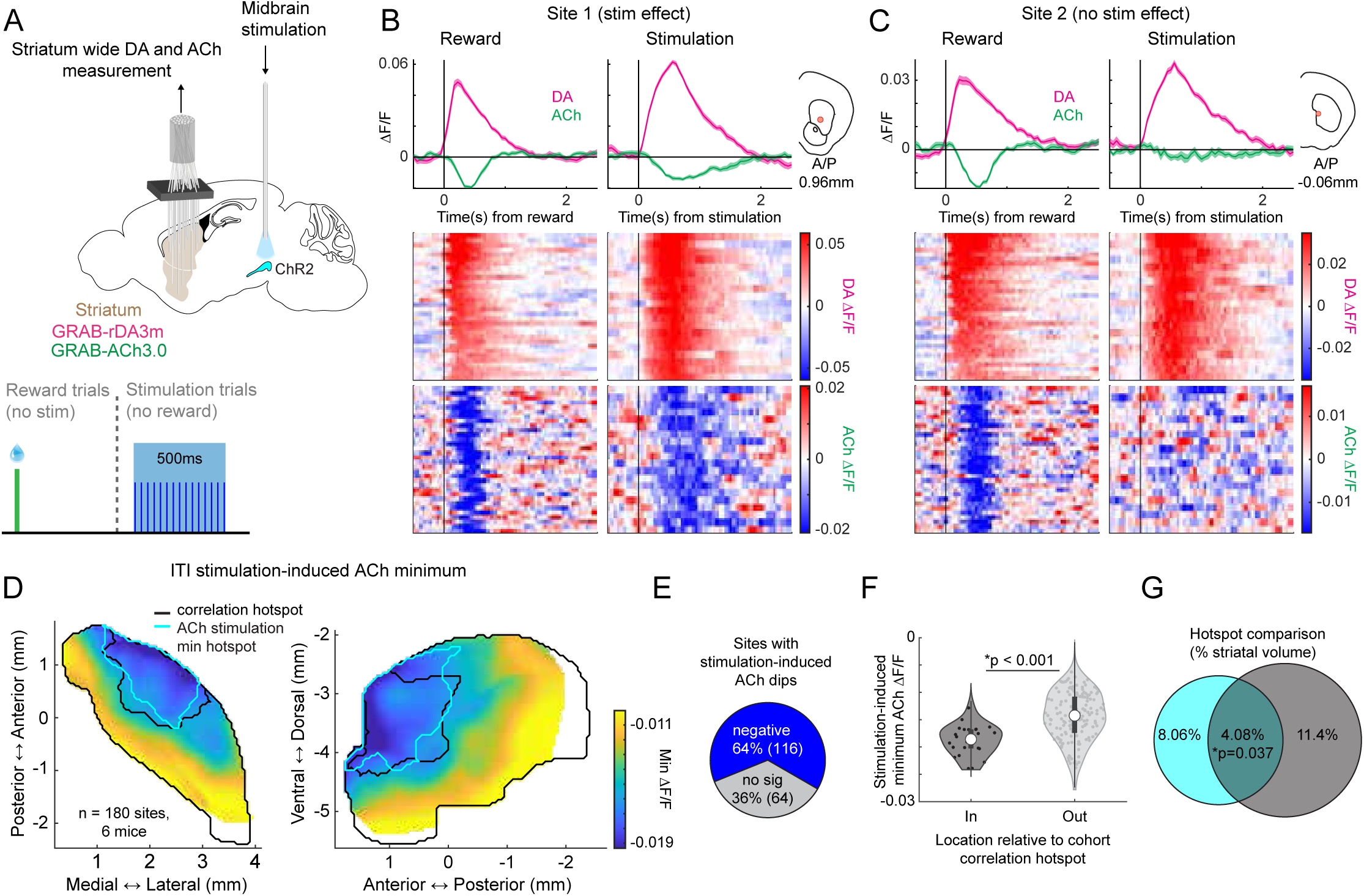
Optogenetic stimulation of midbrain DA neurons is sufficient to evoke ACh dips, selectively in the aDLS hotspot. **a,** Schematic of the experimental paradigm for midbrain optogenetic DA stimulation and simultaneous striatum-wide DA-ACh measurements. **b**, Mean (top) and trial-by-trial (bottom) DA and ACh ΔF/F aligned to unpredicted reward delivery (left) and to DA stimulation onset for a site with a significant stimulation induced ACh dip. Stimulation evoked DA ΔF/F was calibrated to approximately match endogenous reward evoked signals. Inset indicates the site location in the coronal plane. Shaded regions, S.E.M. **c,** Same as b for a site in the same mouse with no significant stimulation effect, despite a reward evoked ACh dip. **d,** Smoothed maps (axial, left; sagittal, right) of average DA stimulation-evoked ACh ΔF/F minima across striatum sites (180 sites, 6 mice). Black contours delineate the cohort cross-correlation hotspot (Ext. Data Fig. 9) and cyan contours delineate the significant hotspot for the stimulation-evoked ACh dips. **e,** Fraction of total sites with significant stimulation-evoked ACh dips. **f,** Violin plots comparing the stimulation-evoked ACh ΔF/F minima for sites inside or outside the aDLS cohort replication correlation hotspot (Ext. Data Fig. 9). Each dot is one site; white circles, median; thick bar, interquartile range; thin lines, 1.5x interquartile range. P-value, Wilcoxon rank-sum test. **g,** Venn diagrams showing the spatial overlap between the global cross-correlation hotspot (black contour, d and Ext. Data Fig. 9) and the ACh minima hotspot (cyan contours in d). Percentages indicate the fraction of the total single-hemisphere striatum volume occupied by individual hotspots (cyan left, black right) and for the intersection of the two hotspots (middle, volume overlap test, see Methods).

### Differential D2R-mediated inhibition across striatal regions inside and outside the aDLS hotspot

Prior work has shown that dopamine (DA) can suppress cholinergic interneuron (CIN) excitability and population synchrony via D2 receptors (D2Rs) in a region-dependent manner ^7,17,39,63^, raising the possibility that spatial differences in D2-mediated inhibition underlie the selective DA-ACh coupling and DA stimulation effects observed *in vivo*. To test this, we compared D2R-dependent regulation of ACh release in acute striatal slices from a region inside the aDLS hotspot and a region outside the hotspot, the nucleus accumbens core (NAcC). Electrically stimulated ACh (5 pulses, 2Hz trains) was measured using the fluorescent sensor GRAB-ACh3.0 in the presence or absence of the D2R agonist quinpirole. Quinpirole suppressed evoked ACh release in both DLS and NAcC (Fig. 8a,b), but the magnitude of suppression was significantly greater in DLS across all stimulation pulses (Fig. 8b).

**Figure 8:**
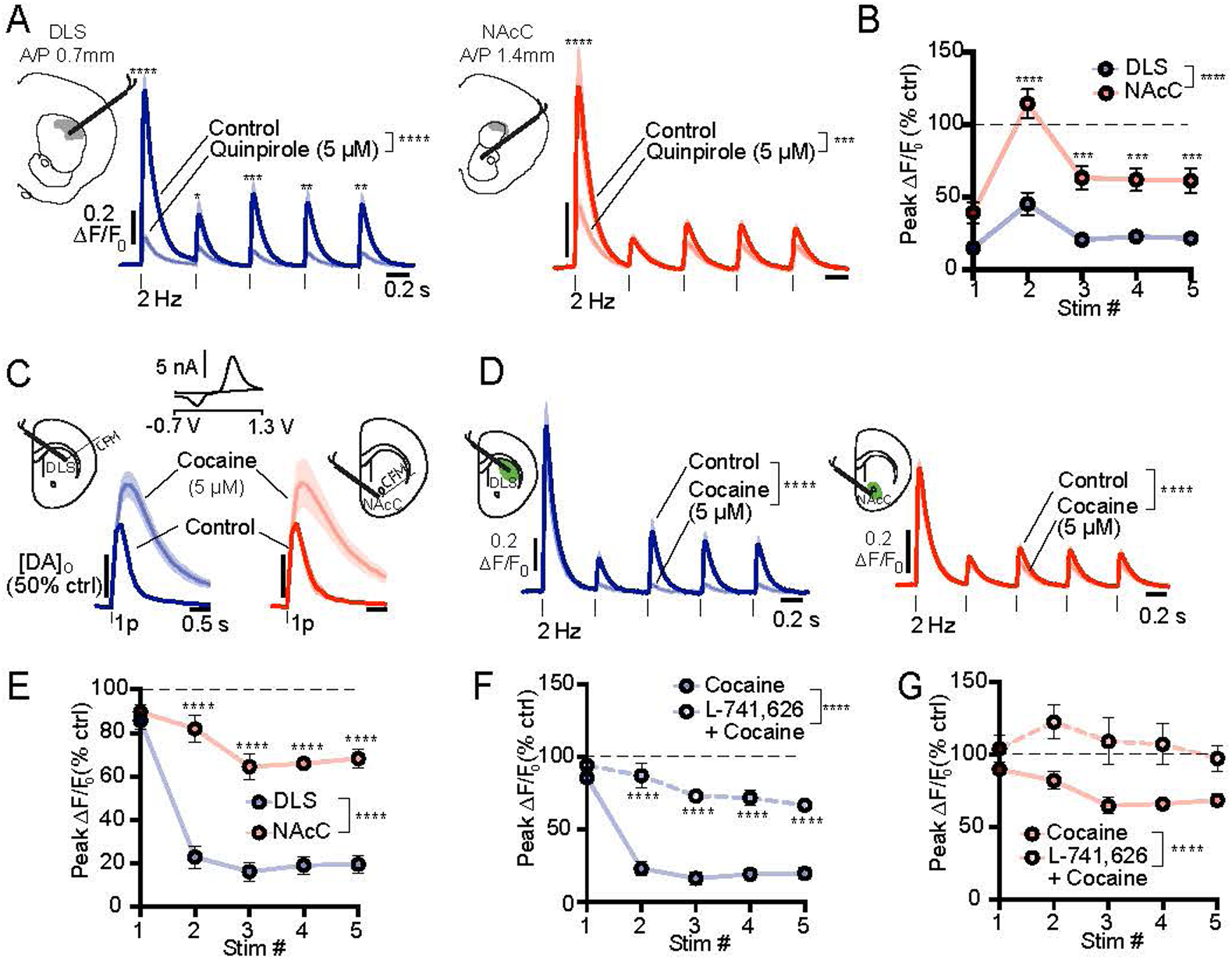
D2-mediated suppression of evoked ACh release *ex-vivo* is stronger in the aDLS hotspot compared to the NACc. **a,** Mean ± SEM ACh ΔF/F versus time evoked by 5 electrical stimulation pulses at 2Hz in DLS (left) and NACc (right) under control conditions (dark) and with the D2R-agonist quinpirole (light), normalized to the first pulse in control. Insets: schematics showing the location of stimulation and imaging in acute coronal slices relative to the aDLS DA-ACh anti-correlation hotspot *in-vivo* (shaded gray region, Fig. 1f). Statistics compare peak ΔF/F for each stimulation in the control and drug conditions. **b,** Mean peak ΔF/F (± SEM) in quinpirole condition normalized to each stimulus in control conditions for DLS (Blue) and NACc (red). Two-way ANOVA, region × stimulus number interaction, F(4,40) = 3.014, p = 0.029; region effect, F(1,40) = 105.9, p < 0.0001; stimulus number effect, F(4,40) = 17.80, p < 0.0001. ***P < 0.001, Bonferroni post-hoc test. n = 5 slices per region from 5 mice. **c,** Cyclic voltammogram and schematic of recording sites (insets). Mean [DA_0_] (± SEM) evoked by single electrical pulses in control and in presence of cocaine in DLS (control, dark blue; cocaine, light blue) and NAcC (control, dark red; cocaine, light red). Data are normalized to single pulse control for each region. Cocaine enhances and prolongs DA signals. Two-way ANOVA: cocaine × region interaction, F(8,36) = 0.20, p = 0.98; cocaine effect, F(8,36) = 5.83, p < 0.0001; region effect, F(1,36) = 0.17, p = 0.67. n = 3 slices from 3 mice. **d,** Mean ± SEM ΔF/F versus time evoked by 5 pulses at 2 Hz in DLS (left) and NACc (right) under control conditions (dark blue) and cocaine application (light blue), normalized to first pulse in control. Insets: schematic of recording region. Statistics compare peak ΔF/F for each stimulation in control and drug condition. **e,** Mean peak ΔF/F (± SEM) in cocaine normalized to each stimulus in control conditions for DLS (blue) and NAcC (red). Two-way ANOVA: stimulus number × region interaction, F(4,55) = 10.75, p < 0.0001; stimulus number effect, F(4,55) = 36.37, p < 0.0001; region effect, F(1,55) = 199.9, p < 0.0001. ***P < 0.0001, **P < 0.01, *P < 0.05, Bonferroni post-hoc test. **f,** Mean peak ΔF/F (± SEM) for each stimulus in cocaine (filled circles) and cocaine in the presence of D2 Blocker L-741,626 (open circles), normalized to control for each stimulus in DLS. Two-way ANOVA: stimulus number × D2R block interaction, F(4,55) = 10.38, p < 0.0001; stimulus number effect, F(4,55) = 34.46, p < 0.0001; D2R block effect, F(1,55) = 231.9, p < 0.0001. **g.** Same as (f) but for NAcC. Two-way ANOVA: stimulus number × D2R block interaction, F(4,40) = 0.87, p = 0.48; stimulus number effect, F(4,40) = 1.51, p = 0.21; D2R block effect, F(1,40) = 32.57, p < 0.0001.

To determine whether elevated endogenous DA would differentially modulate ACh release across regions, we enhanced DA availability with the reuptake inhibitor cocaine, which elevates DA release by similar degree in both regions (Fig. 8c). Consistent with the agonist results, cocaine robustly reduced ACh release at the second through fourth pulses in DLS, while producing only modest suppression in NAcC (Fig. 8d,e). This effect was significantly attenuated by the D2R antagonist L-741,626 in both regions (Fig. 8f,g), confirming that cocaine-induced suppression of ACh release was mediated by increased D2R activation rather than off-target effects. Together, these findings demonstrate that D2R signaling suppresses evoked ACh release more potently in DLS than in NAcC, providing a mechanistic basis for the spatially concentrated DA–ACh coupling and region-selective DA stimulation effects observed *in vivo*.

## Discussion

Here we used large-scale optical fiber arrays to map how endogenous dopamine (DA) and acetylcholine (ACh) release dynamics are coordinated across striatal space and time during behavior. Contrary to the prevailing assumption of homogeneous DA-ACh interactions, we find a pronounced spatial organization in which anti-correlations are strongest within a discrete region of the anterior dorsolateral striatum (aDLS). This was the result of transient DA peaks and dips, followed by lagging ACh dips and peaks, respectively, with magnitudes that were strongly anti-correlated across events. This spatially and temporally structured coupling was consistently expressed across distinct signaling regimes associated with spontaneous locomotion, Pavlovian conditioning, and extinction. In contrast, positive DA–ACh correlations exhibited the opposite temporal ordering (ACh→DA) and lacked a stable spatial organization. Together, these results indicate that DA-ACh coordination is regionally specialized rather than globally conserved and shapes dynamics relevant for movement and learning.

Several lines of evidence indicate that D2 receptor-mediated inhibition of CINs contributes to the aDLS concentrated anti-correlated dynamics. First, negative correlations between DA peaks and lagging ACh release in the aDLS were strongest at latencies consistent with known D2-CIN modulation (>150ms^7,8,10^). Second, optogenetic stimulation of DA neurons alone elicited ACh dips, selectively in the aDLS, at similar latencies, independent of behavior (Fig. 7). Third, *in vitro* recordings revealed stronger D2-dependent inhibition of ACh release in aDLS compared to ventral striatum (Fig. 8). These results, along with prior reports of maximal D2-mediated inhibitory currents in CINs within anterior dorsal striatum^14,17^, indicate that D2 mediated CIN suppression contributes to the concentration of negative DA-ACh interactions in the aDLS. However, this mechanism is unlikely to act in isolation. Short latency ACh dips are likely initiated by feedforward inhibition onto CINs by GABAergic interneurons and inhibitory inputs from the midbrain and globus pallidus^63–69^, and this inhibition is also spatially diverse and modulated by DA^17,63^. Consistent with stronger coupling by feedforward inhibition, we observed a higher probability of DA peaks followed by short-latency ACh dips in the aDLS (Fig. 2b). We therefore propose that the aDLS hotspot for DA-ACh anti-correlations emerges from the convergence of multiple mechanisms acting in concert, including regionally enhanced D2-CINs signaling, feedforward inhibition, and coordinated excitatory inputs across midbrain and striatal circuits.

In addition to dominant anti-correlations, we observed positive DA–ACh correlations at many sites, particularly in posterior, medial, and ventral striatum. These correlations were associated with ACh→DA timing relationships and occurred across behavioral contexts. Although *in vitro* studies demonstrate that ACh can both facilitate and suppress DA release through diverse mechanisms^4,53,55,70–75^, *in vivo* evidence for direct cholinergic control of DA during behavior is mixed ^9,42^. The lack of consistent spatial topography for positive correlations across behavioral epochs (Extended Data Fig. 8e,f) suggests that they may reflect extrinsic, rather than intrinsic, circuit interactions. One parsimonious explanation is that shared, behaviorally evoked glutamatergic input to both DA neurons and CINs produces positively correlated signal magnitudes that vary with task demands. Variation in shared inputs across behavior contexts would yield unstable spatial patterns of positive correlations, which may mask the direct influence of ACh on DA. Isolating direct effects of ACh on DA will require future experiments combining region-specific ACh manipulations with simultaneous DA measurements during defined behaviors.

Consistent with previous work, we observed DA peaks preceding ACh dips to rewards and reward-predictive cues after Pavlovian learning (Figs. 3 and 4^8,42,76^). However, our large-scale recordings revealed that strong trial-by-trial anti-correlations between DA and ACh magnitudes (DA peak vs ACh) were selectively concentrated in the aDLS, despite widespread expression of both signals. This finding supports a model in which DA is neither necessary nor sufficient to generate cue- or reward-evoked ACh dips but instead modulates their magnitude or duration in a region-dependent manner^8,11,17,77^. What is the possible functional consequence of this selective modulation? One possibility is to enhance synchronization of DA peaks and ACh dips to promote more efficient learning related plasticity^5,50,51^. In support of this idea, we have previously found that positive RPE encoding in ACh dips is localized to the aDS, possibly inherited from DA^51,62,78,79^. A second, potentially more critical consequence, emerges during extinction. DA dips to omitted rewards may reduce D2-mediated CIN inhibition, disinhibiting ACh release and permitting transient ACh late peaks^62^. ACh peaks may weaken D1-direct spiny projection neuron synapses (dSPNs) and/or strengthen D2-indirect spiny projection neuron synapses (iSPNs)^80–82^. to down-shift learned actions and cue-values and promote behavioral flexibility^62,83,84^. Our findings suggest that this inversion of negative DA RPEs into ACh excitation is most robust in the aDLS, positioning this region as a key locus for updating behavior in response to negative outcomes.

The importance of DA and ACh interactions for movement control is rooted in the known pathophysiology of Parkinson’s Disease (PD). Loss of substantia nigra DA neurons is thought to disinhibit CINs, producing elevated ACh signaling that contributes to motor symptoms^29–37^. Our results provide insight into how and where DA and ACh signals are expressed and interact during spontaneous locomotion, perhaps shedding light on regional signal disruptions in PD. Previous studies have reported widely varying movement initiation correlates in DA and ACh signals, including increases and decreases^6,40–42,44,45,47^. We mapped a new spatial topography of DA and ACh signals associated with the initiation and invigoration of spontaneous locomotion which may reconcile some of these disparate findings (Fig. 5). Movement initiations were marked by aDLS localized DA dips, followed by more widespread (but aDLS concentrated) ACh peaks. Following initiation, when animals invigorated high velocity locomotion, an opposite sequence was expressed: DA peaks were followed by ACh dips. These results suggest that conflicting prior reports may be due, in part, to where in the striatum or DA midbrain recordings were performed and what phase and type of behavior was examined. Regardless of whether these signals influence behavior on rapid (∼100s of ms) or extended timescales (minutes-days), our data indicate that region-specific DA-ACh dynamics differentially modulate the immediate or long term expression of distinct locomotion phases. Consistent with other behavioral epochs, DA-ACh anti-correlations were concentrated in the aDLS for both movement phases (Fig. 6). This result suggests that DA loss in the PD state may produce region-specific, not homogenous, elevations in phase specific ACh movement signals, suggesting new spatial and temporal targets for future PD therapeutic interventions.

## Methods

### Mice

For in-vivo experiments, we used a total of 17 adult female and male mice (postnatal 3-5 months, weighing 20-30 g): wild-type C57BL6/J (n=11, Jackson Labs, strain #00664) and heterozygous DAT^IREScre^ (n=6, Jackson Labs, strain #006660). Wild-type mice were used for simultaneous DA and ACh extracellular recordings (n=5; 2F, 3M) for spontaneous behavior experiments; n=3 (1F, 2M) for Pavlovian experiments; and n=3 (1F, 2M) for control experiments with non-functional sensors). Heterozygous DAT^IREScre^ mice (n=6; 4F, 2M) were used for midbrain optogenetic stimulation experiments with simultaneous DA and ACh extracellular recordings. Mice were initially group-housed and later housed individually after surgery, under standard laboratory conditions (20-26°C, 30-70% humidity) on a reverse 12-hour light/dark cycle (lights on at 9 p.m.), with free access to food and water, except during water scheduling. All experiments were conducted during the dark cycle. Animal care and procedures were performed in accordance with protocols approved by the Boston University Institutional Animal Care and Use Committee (protocol no. 201800554) and followed to the Guide for Animal Care and Use of Laboratory Animals.

For ex-vivo experiments, mice were group-housed and maintained on a 12- hour light/dark cycle at 22°C and 60-70% humidity. *Ad libitum* access to food and water was provided. All the procedures were carried out with ethical approval from the University of Oxford and under the authority of a UK home office Project License (P9371BF54). Eight WT mice (6M, 2F; Charles River) were used and 5 of them (3M, 2F) were injected with the GRAB ACh sensor at 6-7 weeks and used for experiments 3-5 weeks post-injection. The other three male mice were used for fast-scan cyclic voltammetry experiments.

### Stereotaxic viral injection surgeries for multifiber photometry recordings and optical stimulation of dopamine neurons

Mice were anesthetized with 3-4% isoflurane and positioned in a stereotaxic frame (Kopf instruments) on an electric heating pad (Physitemp instruments). Pre-operative analgesia was administered using extended-release buprenorphine (3.25 mg/kg, subcutaneous, Ethiqa XR). After induction, isoflurane was maintained at 1-2% (in 0.8-1 L min-1 pure oxygen), with body temperature maintained at 37°C throughout surgery. The scalp was shaved and cleaned with iodine solution before performing a large craniotomy over the right hemisphere with a surgical drill (Midwest Tradition 790044, Avtec Dental RMWT), exposing the brain surface from 2.2 to -1.8 mm in the anterior-posterior (AP) direction and 0.49 to 3.4 mm in the medial-lateral (ML) direction relative to bregma. For simultaneous dopamine-acetylcholine extracellular recordings, 350 nl of a 1:6 mix of pAAV-hSyn-rDA3m (5.83 x 10^12^ VG ml^-1^, Bio hippo^59^) and AAV9-hSyn-ACh3.0 (5.86 x 10^12^ GC ml^−1^, Bio hippo ^60^) was pressure-injected into the striatum of WT mice at a rate of 100nl/ min through a pulled glass pipette (tip diameter 30-50 μm) at approximately 30 locations chosen to maximize expression around fiber tips. For control experiments, 500 nl of a 1:67 mix of pAAV-CAG-tdTomato (Adgene #59462) and AAV9-hysn-ACh3.0-mut (WZ Biosciences) ^60^, was injected into the striatum of WT mice using the same strategy.

Following viral injections, the multifiber array was carefully lowered into position and secured with Metabond. The craniotomy was sealed with Kwik-Sil and a metal headplate was then attached for head fixation, and the implant surface was coated with Metabond and carbon powder to reduce optical artifacts. Post-surgically, mice were housed individually with heating pad and received subcutaneous injections of meloxicam (5 mg/kg, Covertus) and 1 mL of saline daily for 4 days. The mice were allowed to recover in their cages for at least 2 weeks before experimental procedures. Multifiber array fabrication has been previously described in detail^58^. For midbrain optogenetic stimulation experiments with simultaneous striatum dopamine-acetylcholine recordings, DAT^IRES^cre received striatal injections of DA and ACh fluorescent sensors and multifiber array implantation following the protocol described above. Three weeks post-injections and implant, these mice underwent a second surgery in which three small craniotomies (1mm diameter) were drilled through the Metabond and skull over the midbrain at pre-marked coordinates. Then pAAV-EF1a-doublefloxed-hChR2(H134R)-EYFP-WPRE-HGHpA (Adgene, # 20298), 2.5 x 10^13^ GCml^-1^ diluted 1:2 in PBS was injected into the right SNc and VTA at 3 sites (200 nl/site at a rate of 100nl/min) at the following three coordinates in mm, AP:-3.05, ML: ± 0.6, DV: -4.25; AP:-3.15, ML: ± 1.3, DV: -4.1; AP:-3.5, ML: ± 1.15, DV: -4.1. Following the injections, a 90 degree 100 µm core mono fiber-optic cannula (MFC_100/125 - 0.66 NA) attached to a zirconia ferrule (Doric) was slowly lowered to a depth of -4 mm from the dura at the coordinates in mm, AP: -3.15 mm and ML: ±1.3 mm. The craniotomies were sealed with Kwik-Sil (WPI) and the optical fibers were secured to the skull with Metabond (Parkell). Mice received the same post-operative care described above and were allowed to recover in their cages for at least one-week post-surgeries.

### Multifiber dual-color photometry recordings and preprocessing

Optical measurement set-up was previously described in detail^58^. Imaging data was acquired using HCImage live (HCImage live, Hamamatsu). Dual wavelength excitation was performed in a quasi-simultaneous externally triggered imaging mode, where the two LEDs were alternated and synchronized with imaging acquisition via 5V digital TTL pulses. 470 nm excitation was alternated with 570 nm excitation at 36 Hz (20 ms exposure time) to achieve a frame rate of 18 Hz for each excitation wavelength. A custom MATLAB software controlled the timing and duration of TTL pulses through a programmable digital acquisition card (NIDAQ, National Instruments PCIe 6343). Voltage pulses were transmitted to the NIDAQ from the camera following the exposure of each frame to confirm proper camera triggering and to synchronize imaging data with behavior data. Videos of the fiber bundle surface were motion-corrected using a previously described whole-frame cross-correlation algorithm ^85,86^ and then processed to extract fluorescence changes from each fiber as previously described^58^. Mean fluorescence time series were extracted from regions of interest (∼25um diameter) defined within each fiber and then normalized to a baseline, defined by fitting a 2-term exponential function to the mean fluorescence time series. To remove low frequency artifacts, the ΔF/F signals were high-pass filtered using a finite impulse response filter with a passband frequency set at 0.3 Hz for green sensors, and 0.1Hz for red sensors. The data in the midbrain DA stimulation experiment (Fig. 7) were not high-pass filtered in order to preserve the longer timescale dynamics of the stimulation effects on ACh.

### Behavior training and recording paradigm

Three weeks post-surgery, mice were placed on a water restriction schedule (1 ml/day) and maintained at 80-85% of their baseline body weight. They were habituated for 3 to 4 days to head-fixation on a lightweight cylindrical running wheel (19 cm diameter, 12 cm width) until they freely ran and transitioned spontaneously between resting and running. The wheel’s linear velocity was sampled at 2kHz by a rotary encoder (E2-5000, US Digital) attached to the wheel axle. During the habituation period, unpredicted 5 µL water rewards (inter-trial interval: 10-30 s) were dispensed through a spout controlled by an electronically controlled solenoid valve, and licking was monitored using a capacitive touch circuit and video recordings. During photometry recordings, mice underwent two distinct daily sessions. In the first session, mice ran and rested on the wheel with no lick spout present and no reward available, allowing for recording of purely spontaneous behavior. In the second session, mice had access to the lick spout and received 5ul unpredicted water rewards delivered at random intervals (10-30s).

### Midbrain optogenetic stimulation

For unilateral optogenetic stimulation of DA neurons during behavior, the implanted optical fibers were connected to a 100 µm patch cord (Doric) with zirconia sleeves and connected to a 450nm laser diode light source (Doric). Laser power at the tip of the optical fibers was 2.5 mW. To activate dopamine neurons, 4 ms blue light pulses were delivered at 30 Hz for 500ms, 100 ms after reward delivery. Additionally, during the ITI, mice received 5 stimulation trains with identical parameters delivered at randomized intervals.

### Pavlovian conditioning and partial extinction

The behavior set up and pavlovian conditioning and extinction paradigm were previously described in detail^62^. Briefly, mice were water restricted and acclimated to head fixation on a spherical Styrofoam treadmill for 3 to 4 days before the Pavlovian conditioning and photometry recordings. During experiments, mice were free to move on the treadmill while ball rotation was measured using optical mice and an acquisition board. Three weeks after multifiber fiber array implantation, mice began a dual cue delay Pavlovian conditioning task. Each day, mice received 60 cue presentations (30 light and 30 tone) in a pseudorandom order. Light cues, delivered via a 7mW LED positioned 20 cm from the mouse, and tone cues (12 kHz, 80 dB) presented through a USB speaker 30 cm away, each lasted 6 seconds and were followed by water reward after a 3-second delay. Inter-trial intervals (ITIs) ranged from 4 to 40 seconds, with eight additional non-contingent rewards per session. Training continued for 26-38 days until mice learned to associate both cues with rewards. Subsequently, mice underwent a partial extinction phase in which the reward probability for one cue was downshifted to 20% while the other cue to 80%. Training persisted until significant reductions in responses to the 20% cue were observed. Cue probabilities were then reversed, with the order counterbalanced across mice.

### Brain preparation for CT scan and Immunohistology

At the end of recordings and behavioral experiments, mice were administered Euthasol (400 mg/kg, Covertus Euthanaisa III) via intraperitoneal injection and then perfused intracardially with phosphate-buffered saline (PBS 1%, Fisher Scientific). Following decapitation, the ventral skull was removed to expose the ventral brain surface to facilitate the penetration of contrast agent (Lugol, Carolina Scientific). Implanted brains were maintained intact and post-fixed in 4% paraformaldehyde for 24 hours, rinsed with 1% PBS, and transferred to a 1:3 dilution of lugol solution in distilled water for 4 to 5 days. Lugol solution diffusion into brain tissue enhances contrast for CT imaging, enabling fiber localization. CT scanning and fiber localization procedures were previously described in detail^58^. Following CT imaging, Lugol solution was cleared from brain tissue by immersing the brains in 5% sodium thiosulfate (STS, Carolina Scientific) dissolved in 1% PBS for 4-6 days. After implant extraction, brains were returned to the STS solution for one hour before transfer to 30% sucrose in PBS 1%. Once brains equilibrated and sank, 50 µm coronal sections were prepared using a cryostat (Leica CM3050 S) and stored in PBS 1%. For immunostaining of ChR2 expression, free-floating section were washed three times for 10 min in PBS, then incubated in PBS containing 0.2% Triton X-100 and 5% normal goat serum (NGS) for 2 hours at room temperature to block nonspecific binding and permeabilize the cells membrane. Sections were then incubated with primary antibody (chicken anti-GFP, 1:1000, ThermoFisher Scientific, #A10262) in PBST with 5% NGS for 48 hours at 4° C on a shaker. Following three 10-minute washes in PBS, sections were incubated with secondary antibody (Alexa Fluor 488 goat anti-chicken, 1:200, ThermoFisher, A11039) in PBST with 5% NGS for 2 hours at room temperature on a shaker, protected from light. Sections were then washed three times for 10 minutes in PBS and then mounted on slides using Vectashield mounting medium. Confocal microscopic images were acquired on a Zeiss LSM 800.

### Stereotaxic intracranial injections for ex-vivo experiments

Mice were anaesthetized with isoflurane and transferred to a stereotaxic frame (Kopf Instruments). The skull was exposed, craniotomies were performed with a small drill and adeno-associated virus (AAV)-packaged constructs (1 mL/site) were injected at a rate of 200 nL/min with a 32-gauge Hamilton syringe (Hamilton Company), withdrawn 5 min after injection. AAV5-hSyn-ACh3.0 or AAV9-hSyn-ACh3.0 (∼ 0.8 x 1013 genome copies/mL) were bilaterally injected in the dorsolateral striatum (ML ±2.2 mm from bregma, AP +0.7 mm from bregma, DV -2.7 to -2.5 mm from top of the brain) and in the NAcC (ML ±0.9 mm from bregma, AP +1.4 mm from bregma, DV -3.8 to -3.5 mm from top of the brain).

### Slice preparation

Mice were sacrificed by cervical dislocation and subsequently decapitated for collection of the brain. For all imaging and FCV experiments, acute 300 μm thick striatal coronal slices were prepared using a vibratome (VT1200S, Leica) in ice-cold HEPES-based buffer saturated with 95% O2/ 5% CO2, containing (in mM): NaCl (120), NaHCO3 (20), HEPES acid (6.7), KCl (5), HEPES salt (3.3), CaCl2 (2), MgSO4 (2), KH2PO4 (1.2), glucose (10). Slices were maintained in a holding chamber for at least one hour at room temperature (20-22° C) in HEPES-based buffer before transferal to the recording chamber.

### Fast-scan cyclic voltammetry

Slices were bisected and transferred to the recording chamber where they were superfused at ∼ 1.8 ml/min in bicarbonate-buffer based artificial cerebrospinal fluid (aCSF) containing (in mM): NaCl (125), NaHCO3 (26), KCl (3.8), CaCl2 (2.4), MgSO4 (1.3), KH2PO4 (1.2), glucose (10), saturated with 95% O2/5% CO2, at 31-33° C. Slices were left to equilibrate in the bath for 30 minutes while recording electrode was charged in tissue. Evoked extracellular DA concentration ([DA]o) was measured using FCV with custom-made carbon fiber microelectrodes (CFMs) and a Millar voltammeter (Julian Millar, Barts and the London School of Medicine and Dentistry). A triangular voltage waveform (- 700 to +1300 mV range vs Ag/AgCl reference electrode) was scanned across the CFM at a scan rate of 800 V/s and sampling frequency of 8 Hz. Evoked currents were attributed to DA by comparison of the potentials for peak oxidation and reduction with those of DA in aCSF (oxidation peak +500-600 mV, reduction peak -200 mV). Currents sampled at the DA oxidation peak potential were measured in each voltammogram from baseline and plotted against time, to provide profiles of [DA]o versus time. DA was electrically evoked with a surface bipolar concentric Pt/Ir electrode (25/125 μm inner/outer tip diameter, FHC Inc.). Recording electrodes were calibrated in 2 μM DA in each experimental media at the end of each experiment. Calibration solutions were prepared before use from 2.5/5 mM DA in 0.1 M HClO4 stock solution stored at 4°C.

### Ex-vivo GRAB ACh sensor imaging

Slices were bisected and transferred to the recording chamber, where they were superfused at ∼ 2-2.5 ml/min in bicarbonate-buffer based artificial cerebrospinal fluid (aCSF) containing (in mM): NaCl (125), NaHCO3 (26), KCl (3.8), CaCl2 (2.4), MgSO4 (1.3), KH2PO4 (1.2), glucose (10), saturated with 95% O2/5% CO2, at 31-33° C. Slices were left to equilibrate in the bath for 30 min prior recording. Slices were imaged under a 10x immersion objective (Olympus) using a sCMOS camera (IRIS9, Teledyne Photometrics) and an ImageJ plugin (Micro-Manager 1.4). For experiments involving electrical stimulation, the stimulating electrode was positioned in sites with the brightest expression. Electrical stimulation was performed with a surface bipolar concentric Pt/Ir electrode (25/125 μm inner/outer tip diameter, FHC Inc.) and stimuli were applied at 0.6 mA for 200 μs every 2.5 min. Images were acquired at a rate of 100 Hz (camera exposure 10 ms) under continuous blue light (410 nm LED, 5-10 mW intensity, 2-3.5 s duration) and camera pixels were binned 2 x 2. Electrical stimulation, LED light and image acquisition were synchronized using TTL-driven stimuli via Multi Channel Stimulus II (Multi channel systems).

### Drugs

All drugs for ex vivo experiment were prepared as follows: Quinpirole (5µM, Cambridge Bioscience), L-741,626 (1µM, Abcam) and cocaine (5µM, Sigma-Aldrich) were diluted to 1000x stock aliquots and stored at -20°C. Final drugs concentrations were prepared in aCSF saturated with 95% O_2_ /5% CO_2_ immediately before use.

### Quantification and statistical analysis for in vivo data Cross-Correlation Analyses

Spontaneous cross-correlation at each site was calculated as the Pearson correlation of DA vs ACh shifted at lags within +/-1s with respect to DA, generating 37 correlation coefficients (18Hz sampling rate). We present the data with respect to DA such that a positive latency (or lag) indicates DA-leading correlation, and a negative latency indicates a ACh-leading correlation. Dominant correlation coefficient was the min or max Pearson’s *r*, whichever had the larger magnitude. For the sliding correlation analysis (Ext. Fig. 3a-c), correlation was calculated over a sliding 2-second window (width of 36 samples, sliding step of 1 sample) at the time-lag of dominant correlation (Fig. 1e).

Event-related cross-correlation analyses were calculated correlating the magnitude of the trial-by-trial event of interest, e.g., DA peaks, with trial-by-trial magnitude of the other neuromodulator, e.g., ACh, at different time points within the window of +/-500ms with respect to the event of interest. For example, for the unpredicted reward analysis, trial-by-trial DA peak magnitude was correlated with trial-by-trial ACh at t=-500ms,… t=0ms, … and then t=500ms with respect to the time of the DA peak to generate 19 correlation coefficients (sampling rate = 18Hz) for each site. The dominant correlation coefficient was the min or max Pearson’s *r*, whichever had larger magnitude. For illustrative purposes, sites were then clustered into 2 clusters using k-means based on correlation coefficient and corresponding latency.

### Smooth volumetric maps

Smooth volumetric maps with were generated via 3-D natural neighbor interpolation using Matlab’s *scatteredInterpolant* with 50-micron isotropic voxel size, smoothed with a 3-D Gaussian smoothing kernel with standard deviation of 2 voxels, and then masked with the striatum volume mask generated from the Allen Brain Common Coordinate Framework Mouse Atlas. Maps shown in 2-D are mean projections.

### Hotspot identification and analyses

Hotspots were defined according to 2 criteria: values had to be significantly concentrated in that region (as opposed to randomly dispersed) and the mean calculated over the region had to exceed the mean calculated over the whole striatum. Significant concentration was assessed via local spatial autocorrelation, calculated for each voxel using Local Moran’s *I*, a Local Indicator of Spatial Association (LISA)^61^, over a neighborhood defined via queen’s weight matrix of +/-500microns. This neighborhood was chosen based on inter-fiber distances to include at least 3 fibers. A bootstrapped null distribution (n = 10000) was then generated by randomly shuffling the values across recording sites, re-smoothing, and re-calculated Local Moran’s *I*. Significance was determined at each voxel based on an alpha value of 0.05, i.e., exceeding the 95th percentile of the null distribution. Regional hotspot volumes were defined as connected volumes of at least 1000 voxels (corresponding to 500microns^3^, equivalent in volume to the neighborhood definition used for the Local Moran’s *I* calculation) with significant local Moran’s *I* and with mean value (i.e., correlation coefficient) exceeding the the striatal mean in the appropriate direction. For example, for minimum correlation coefficient hotspot, the mean correlation coefficient calculated from the hotspot volume (smoothed map) had to be lower than the mean correlation coefficient calculated over the whole striatum (smoothed map).

To test whether two hotspot volumes A and B significantly overlapped, we devised a volume overlap test. A bootstrapped null distribution of chance overlap was generated by randomly selecting a connected volume within the striatum of the same size as A and a (connected) volume within the striatum of the same size as B, and determining the overlap, repeated 10000 times. The p-value for the overlap was calculated as the percentage of the bootstrapped null distribution overlap values greater than or equal to the actual overlap.

### Transient detection and analysis

We reasoned that true transients have some temporal autocorrelation vs single-timepoint jumps in magnitude, which are more likely noise. Therefore, significant transients were defined as having at least 3 consecutive timepoints above the 97.5th percentile of the null distribution (peaks) or at least 3 consecutive timepoints below the 2.5th percentile of the null distribution (troughs). The null distribution was calculated as the mean of 3 randomly sampled datapoints from the fluorescence timeseries, repeated 10000 times. Transient detection was also used to determine whether sites had signal or not: sites were determined to have signal if the rate of transient detection exceeded chance. For the analysis relating moment-to-moment correlation strength with the presence of DA and ACh transients (Ext. Data Fig. 3a-c), the presence of transients was calculated as the percent of the 2-second time window in which a transient (positive or negative) was present for DA, plus that for ACh.

Paired transients were defined as DA and ACh transients with peaks or troughs occurring within 1s of each other. Note that in the event, for example, that a DA peak has 2 ACh troughs following it, both occurring within the 1s allowance window, only the first ACh trough is counted.

For each site, the directionality index for paired transients was calculated as (A - B) / (A + B), where A was the number of pairs where the ACh transient preceded the DA transient, and B was the number of pairs where the DA transient preceded the ACh transient. A value of -1 indicates that all pairs are DA-leading, and a value of +1 indicates that all pairs are ACh-leading. For each site, pairing occurrence was calculated as the percent of the candidate transients that are paired, concretely (ACh_paired_ + DA_paired_)/(ACh_all_ + DA_all_), where ACh_paired_ is the number of ACh transients that are part of specific pair (e.g., ACh peaks accompanied by DA peaks) and ACh_all_ is the total number of ACh transients (e.g.,all ACh peaks), and similarly for DA.

Null pairing rate was calculated by 10000 repetitions of shuffling DA and ACh traces across days (e.g., the DA trace from one day ends up paired with the ACh trace for a randomly selected different day), such that the general temporal dynamics were preserved but the recordings were non-simultaneous, giving an estimate of the rate of chance detection of pairing.

Paired transient magnitude correlation was calculated by correlating the min (for troughs) or max (for peaks) of DA transients with those of ACh transients across pair occurrences. Since the number of pairs varied across striatal sites (Fig. 2b) and could therefore modulate the strength of the correlation, magnitude correlation was calculated as follows: for each site, 100 pairs were randomly selected (sites with fewer than 100 occurrences of a pair type were excluded) for the correlation analysis. This was repeated 10000 times, and the resulting mean correlation coefficient recorded for each site; the correlation was deemed significant if 0 fell outside of the 2.5th-97.5th percentile or the 10000 correlation coefficients. Note that we correlated *signed* magnitudes, such that a negative correlation between DA peaks and ACh troughs means that higher DA peaks are associated with deeper ACh troughs.

### Locomotion quantification

Velocity was low-pass filtered at 1.3Hz to account for walking footfall frequency^87^ (the value was taken as mean - 2*SD), and then smoothed over a 300ms window. Clean locomotion onsets were defined as a period of stillness of at least 1s, followed by a bout of forward locomotion of at least 2.27s duration. The velocity threshold for stillness that allowed for postural adjustment micro movements was determined by median velocity during reward consumption. The duration threshold for a locomotion bout was determined by fitting a gaussian mixture model (4 components, chosen via elbow method) to the distribution of bout durations across all mice. The threshold of 2.27s is the mean of the 2nd smallest distribution, the smallest distribution determined to be short duration “jerky” movements or initiations not resulting in locomotion bouts.

Locomotion “invigoration” events were defined as the first local max in the acceleration after an initiation onset. Acceleration was calculated as the timepoint-to-timepoint difference in the velocity, and then, like the velocity, was low-pass filtered at 1.3Hz, and smoothed over a 300ms window.

### Generalized linear model for analysis of locomotion invigoration events

Because locomotion invigoration events occurred with such short latency to initiation events (Fig. 5H, bottom), a generalized linear model was used to estimate the separate contributions of these different events to the DA and ACh signal. Continuous velocity and acceleration were included as continuous predictors, and initiation and invigoration events were modeled with a finite impulse response (FIR) basis set spanning -0.5 to 1s around the event. Concretely, we used MATLAB’s *glmfit* to fit the following model:

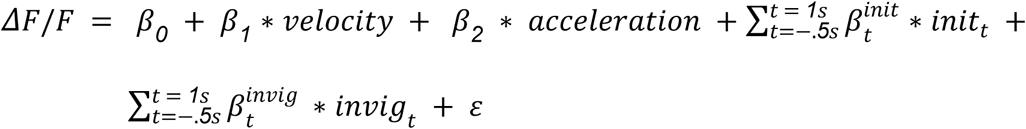

The variable *init_t_* is binary, taking ‘1’ at time *t* with respect to initiation occurrences and ‘0’ elsewhere, and the variable *invig_t_* is binary, taking ‘1’ at time *t* with respect to invigoration events and ‘0’ elsewhere.

The 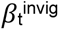 coefficients were used to estimate the overall invigoration-triggered average for DA and ACh. In order to carry out the event-related cross-correlation analysis for invigoration events, trial-by-trial invigoration-triggered activity was estimated using the residuals after regressing out the contributions of velocity, acceleration, and initiation events using the coefficients estimated from the model

### Triggered average significance test

Triggered averages were determined to have significant peaks or dips if, for at least 3 consecutive timepoints, the mean was greater than the 97.5th percentile or less than the 2.5th percentile, respectively, of the bootstrapped null distribution (p<1.525x10^-5^). The bootstrapped null distribution was generated by 10000 iterations of calculating the triggered average from *n* randomly selected events, where *n* = the number of actual number of events contributing to the triggered average of interest.

### Quantification and statistical analysis for ex-vivo data

For experiments, where evoked ACh release was measured with GRAB_ACh_, fluorescence and frame values were extracted from square ROIs (100 x 100 μm) positioned 100 μm away from the tip of the stimulating electrode with a custom-written ImageJ macro. Background was subtracted as previously described^88^. Change in fluorescence from baseline was calculated in MATLAB R2020_a (MathWorks) by fitting a two-term exponential decay function (f(x) = a^bx^ + c^dx^, exp2 function) to pre- and post-peak raw fluorescence values to obtain baseline fluorescence values (F_0_). Fluorescent responses were then calculated as ΔF/F_0_ = (F-F_0_)/F_0_. FCV data were acquired and analysed using Axoscope 10.5 (Molecular Devices) and locally written Microsoft Excel macros (Visual Basic). DA oxidation currents values were measured from background-subtracted voltammograms, converted to concentration using the electrode calibration factor and plotted against time to obtain [DA]o transients. For all experimental sets, data were collected from a minimum of 3 animals. Statistical analysis was carried out using GraphPad Prism (Version 9.4.1) with two-way ANOVA followed by Bonferroni multiple comparisons test. The threshold for statistical significance was set at p < 0.05.

## Acknowledgements

This work was supported by the following funding sources: MWH - Aligning Science Across Parkinson’s (ASAP, ASAP-020370) through the Michael J. Fox Foundation for Parkinson’s Research (MJFF), National Institute of Mental Health (R01 MH125835), Whitehall Foundation Fellowship, Klingenstein-Simons Foundation Fellowship, Parkinson’s Foundation (Stanley Fahn Junior Faculty Award, PF-SF-JFA-836662); MTV - NIMH F32MH120894.

We thank the Boston University Centers for Neurophotonics and Systems Neuroscience for financial and technical support, Micro CT Core, especially Sydney Holder, for providing equipment and technical expertise for micro-CT scanning, and Boston University Animal Science Center for providing central laboratory animal care and support resources. We thank our ASAP team for feedback on the manuscript.

## Data and Code Availability

The data, code, protocols, and key lab materials used and generated in this study will be listed in a Key Resource Table alongside their persistent identifiers at 10.5281/zenodo.18202873

---

**Extended Data Figure 1:**
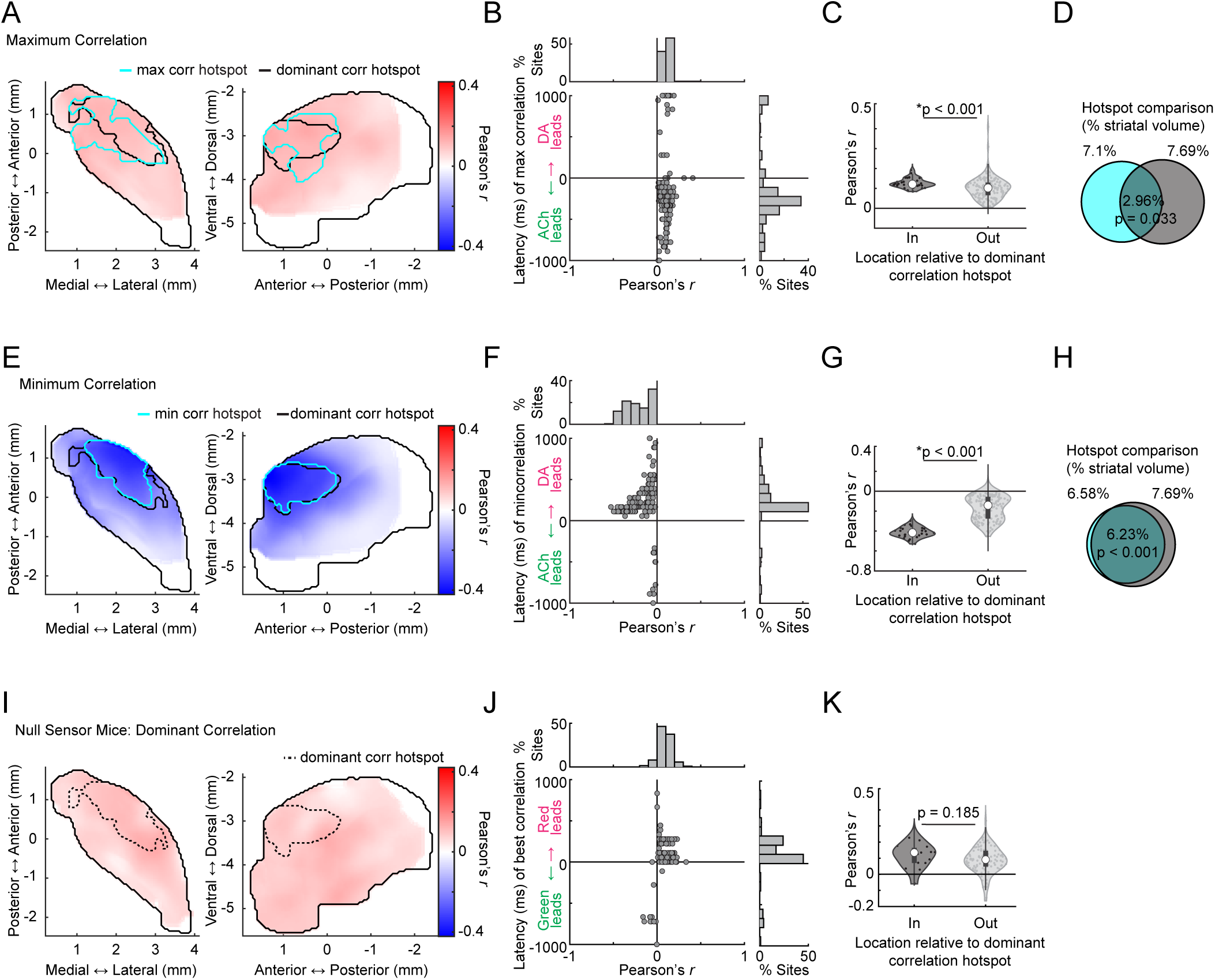
Distributions of cross-correlation maxima and minima and absence of spatially concentrated correlations in control mice. **a,** Smoothed maps (axial, left; sagittal, right) of maximum DA-ACh cross-correlation coefficients across striatum sites. Black contours delineate the dominant cross-correlation hotspot (Fig.1f) and cyan contours delineate significant hotspot for cross-correlation coefficient maxima (1s window). **b,** DA-ACh cross-correlation coefficient maximum vs latency for each site (dot) across all mice (5 mice, 174 sites). Histograms show the percent of total sites binned across correlation coefficients (top) and latencies (right). **c,** Violin plot of DA-ACh correlation coefficient maxima for sites inside and outside the aDLS hotspot. Each dot is one site; white circles, median; thick bar, interquartile range; thin lines, 1.5x interquartile range. Sites inside the hotspot show significantly stronger positive correlations (*p=3.44e-4, Wilcoxon rank-sum test). **d,** Venn diagram showing the spatial overlap between the dominant cross-correlation hotspot (black contour in a) and the maximum correlation hotspot (cyan contour in a, Fig. 1f). Percentages indicate the fraction of the total single-hemisphere striatum volume occupied by each individual hotspot in a (cyan left, black right) and for the intersection of the two hotspots (middle). **e-h,** Same as a-d for cross-correlation minima. **i-k,** Same as a-c for dominant red-green fluorescence correlation coefficients (Fig. 1) in mice (n = 3) expressing the non-functional mutant ACh sensor and static red indicator tdTomato showing sparse weak positive correlations with no anatomical organisation. Dashed contours delineate the cross-correlation hotspot (Fig.1f). No significant hotspot was identified for these mice.

**Extended Data Figure 2:**
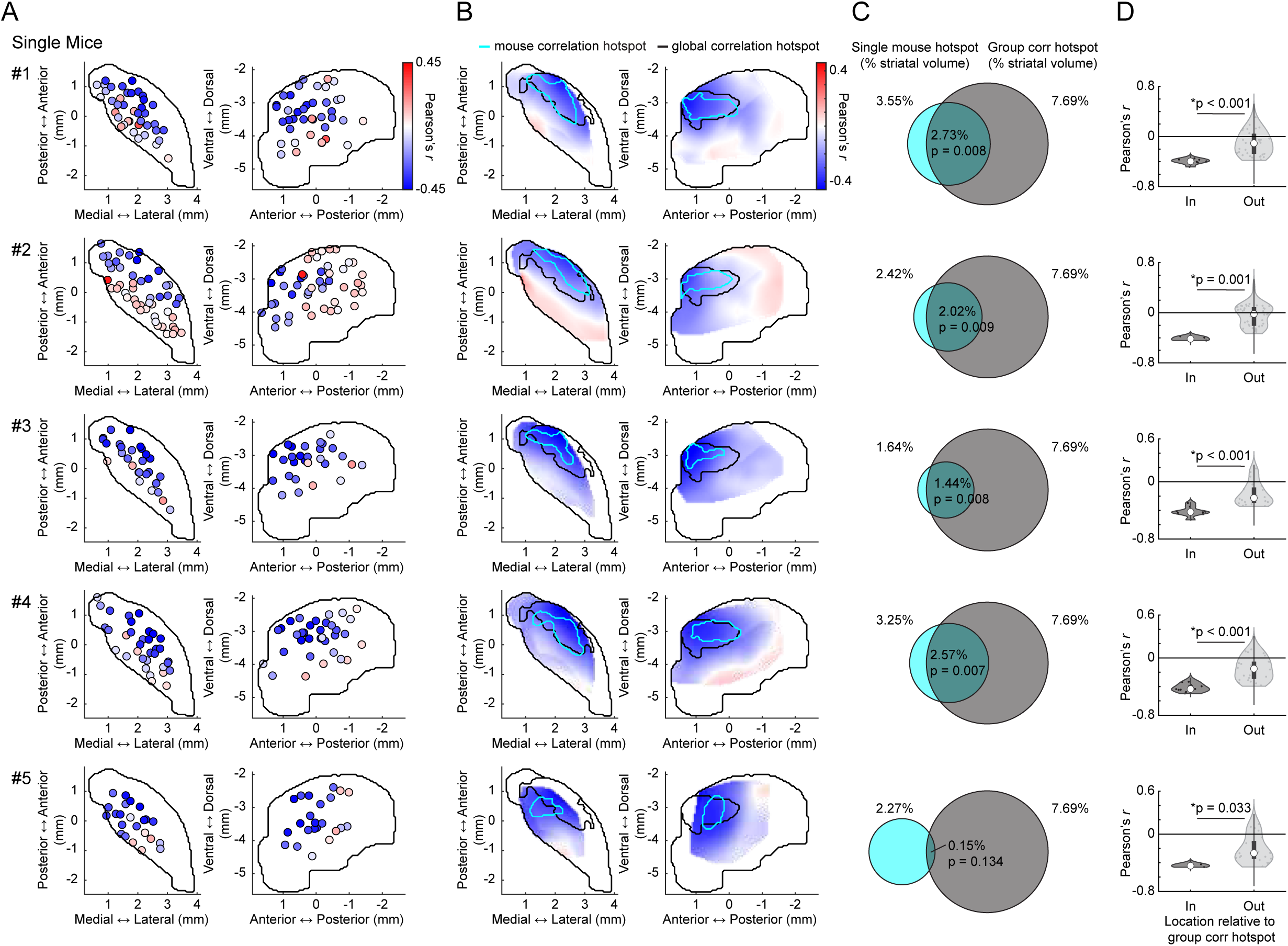
Consistency of the aDLS hotspot for dominant DA-ACh anti-correlations across individual mice. **a,** Maps (axial, left; sagittal, right) of dominant DA-ACh cross-correlation coefficients for individual recording sites (dots) in each mouse. **b,** Spatially smoothed versions of the maps in a. Black contours delineate the global (group) dominant cross-correlation hotspot (Fig.1f) and cyan contours delineate the significant dominant cross-correlation hotspot for each mouse (1s window). **c,** Venn diagrams showing the spatial overlap between the global dominant cross-correlation hotspot (black contour), and the significant hotspot identified in each individual mouse (cyan contour). Percentages indicate the fraction of the total single-hemisphere striatum volume occupied by each individual hotspot in a (cyan left, black right) and for the intersection of the two hotspots (middle). Only one mouse (#5) did not have a dominant correlation hotspot that significantly overlapped with the global hotspot across mice, likely because of incomplete sampling. **d,** Violin plots of DA-ACh dominant correlation coefficients for sites inside and outside the global (group) correlation hotspot for each mouse. Each dot is one site; white circles, median; thick bar, interquartile range; thin lines, 1.5x interquartile range (*p<0.05, Wilcoxon rank-sum test).

**Extended Data Figure 3:**
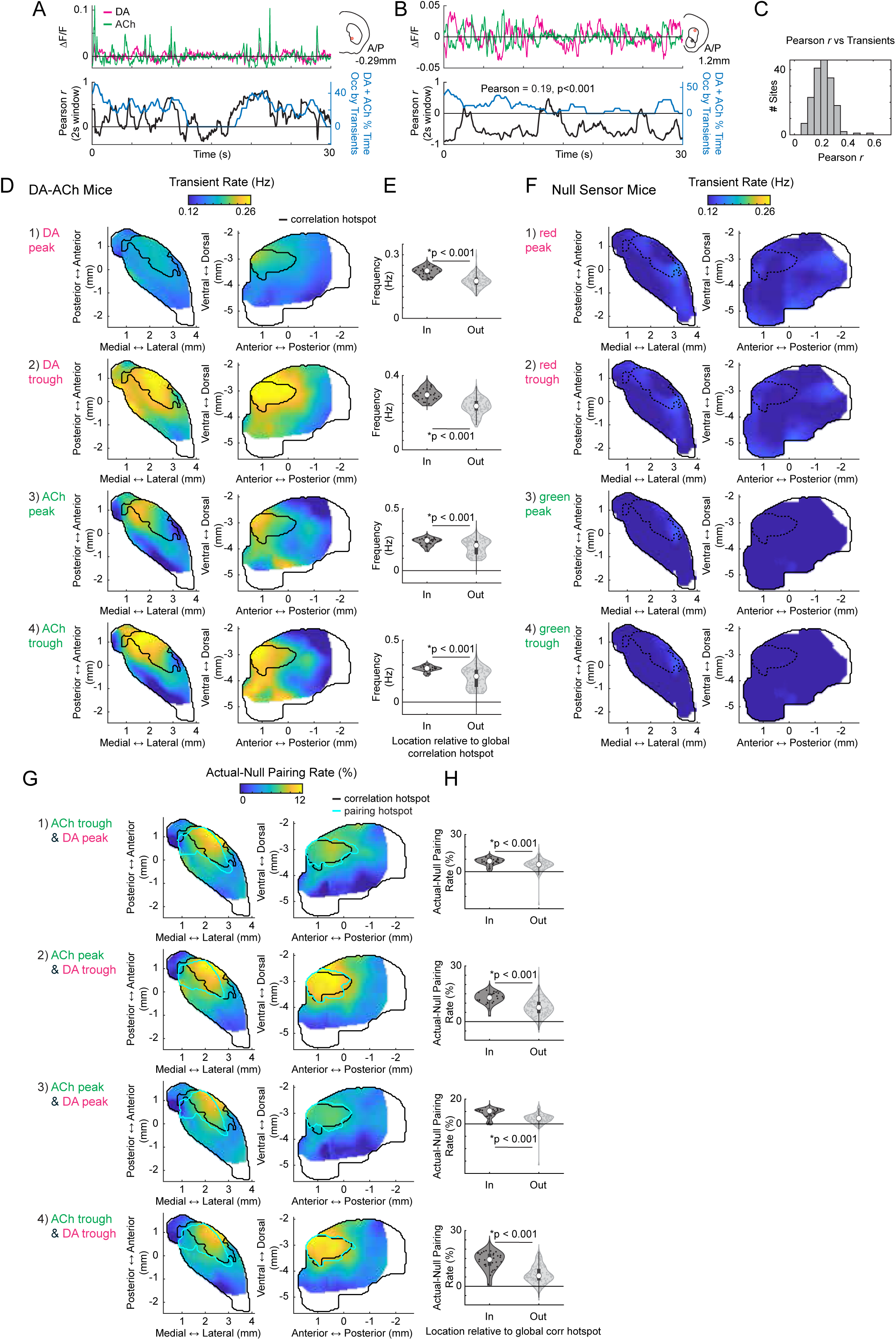
DA-ACh transient frequencies and inverse polarity transient pairing rates are concentrated in the aDLS hotspot. **a,** Top: example DA and ACh ΔF/F traces over a 30s period from a single site with a dominant positive DA-ACh cross-correlation coefficient. Inset indicates the site location in the coronal plane. Bottom: DA-ACh correlation coefficients (Pearson’s *r*) calculated over sliding 2-second windows (black), and the percent time within each window occupied by DA and ACh transients (summed percentages for each, blue). DA-ACh correlation coefficient magnitudes are significantly correlated with the transient prevalence over time (Pearson’s r = 0.47, p < 0.001). **b,** Same as a for another example site with a dominant negative correlation. DA-ACh correlation coefficient magnitudes are significantly correlated with the transient prevalence over time (Pearson’s r = 0.19, p < 0.001). **c,** Histogram of correlation coefficients across sites for correlations between transient prevalence and absolute DA-ACh correlation coefficient magnitudes across all time periods. **d,** Smoothed maps (axial, left; sagittal, right) of transient rates across striatum sites for each DA and ACh transient type (peak or trough). Black contours delineate the global cross-correlation hotspot (Fig. 1f). **e,** Violin plots of transient rates for sites inside and outside the global correlation hotspot for transient types indicated by corresponding rows of d. Each dot is one site; white circles, median; thick bar, interquartile range; thin lines, 1.5x interquartile range. Sites inside the hotspot showed significantly higher transient rates (*p<0.05, Wilcoxon rank-sum test). **f,** Same as d for control mice expressing the null ACh sensor and static tdTomato. False identified transients occurred at low frequency relative to the true sensor and exhibited no significant spatial pattern. Black contours delineate the global cross-correlation hotspot (Fig. 1f). **g,** Smoothed maps (axial, left; sagittal, right) of actual-minus-null occurrence (%) of each set of transient pairs. Black contours delineate the global cross-correlation hotspot (Fig. 1f); cyan contours delineate the hotspots for the actual occurrence of each set of transient pairs from Fig. 2b. **h,** Violin plot of actual-minus-null occurrence of each set of transient pairs for sites inside and outside the global aDLS cross-correlation hotspot. Each dot is one site; white circles, median; thick bar, interquartile range; thin lines, 1.5x interquartile range. Sites inside the hotspot showed significantly higher occurrence of pairs (*p<0.05, Wilcoxon rank-sum test). Pair categories match corresponding rows in g.

**Extended Data Figure 4:**
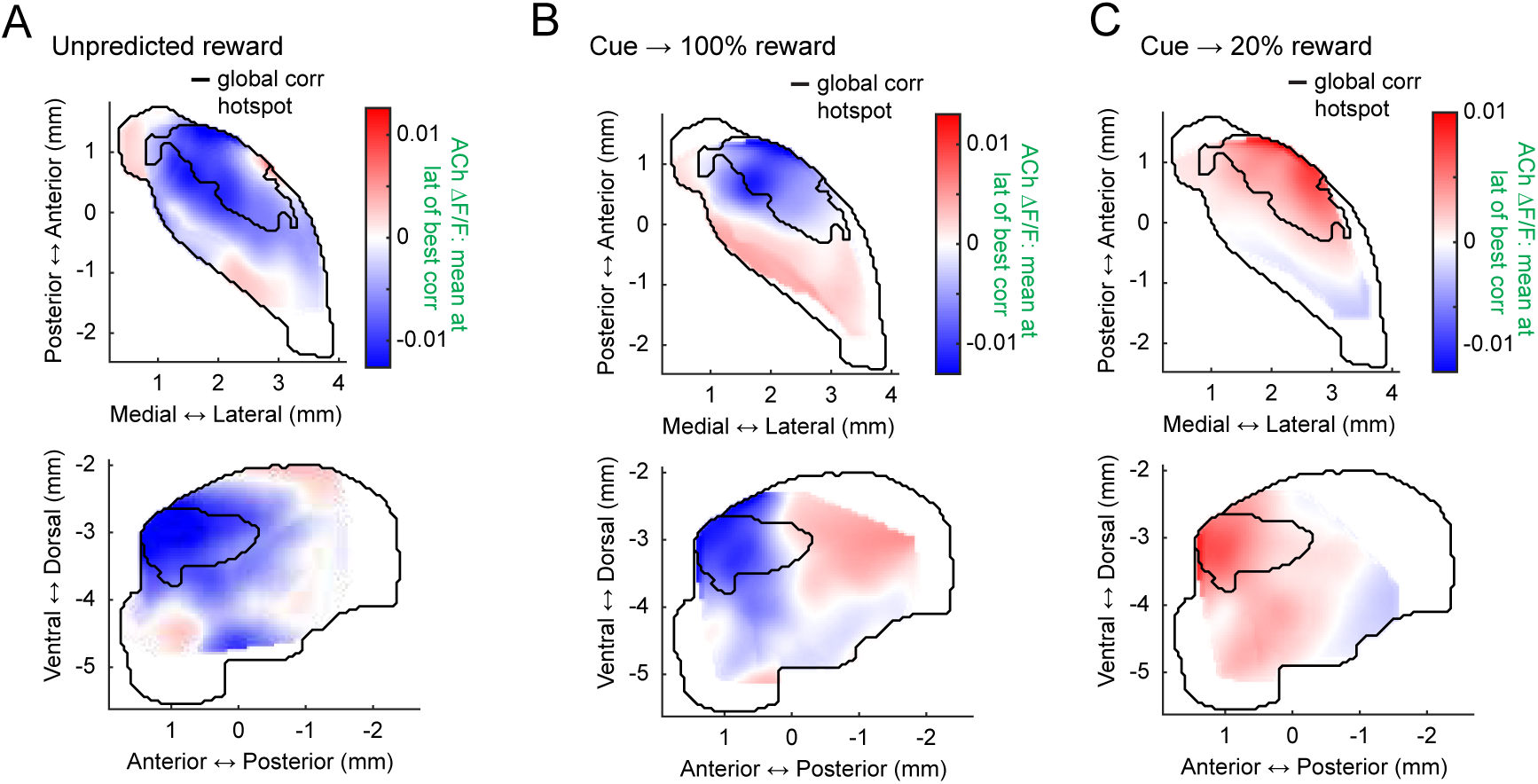
The aDLS anti-correlation hotspot is associated with ACh dips to unpredicted rewards and Pavlovian cues and with peaks to extinguished cues. **a,** Smoothed maps (axial, top; sagittal, bottom) of mean ACh ΔF/F at the latency of the dominant correlation coefficient for unpredicted reward deliveries. The black contour delineates the global cross-correlation hotspot (Fig. 1f). **b,** Same as a for conditioned cues. **c,** Same as a for cues following extinction. Note that mean ACh ΔF/F at the time of peak negative correlations in the aDLS hotspot is positive, corresponding to inverse dynamics relative to initial conditioning (DA dip → ACh peak).

**Extended Data Figure 5:**
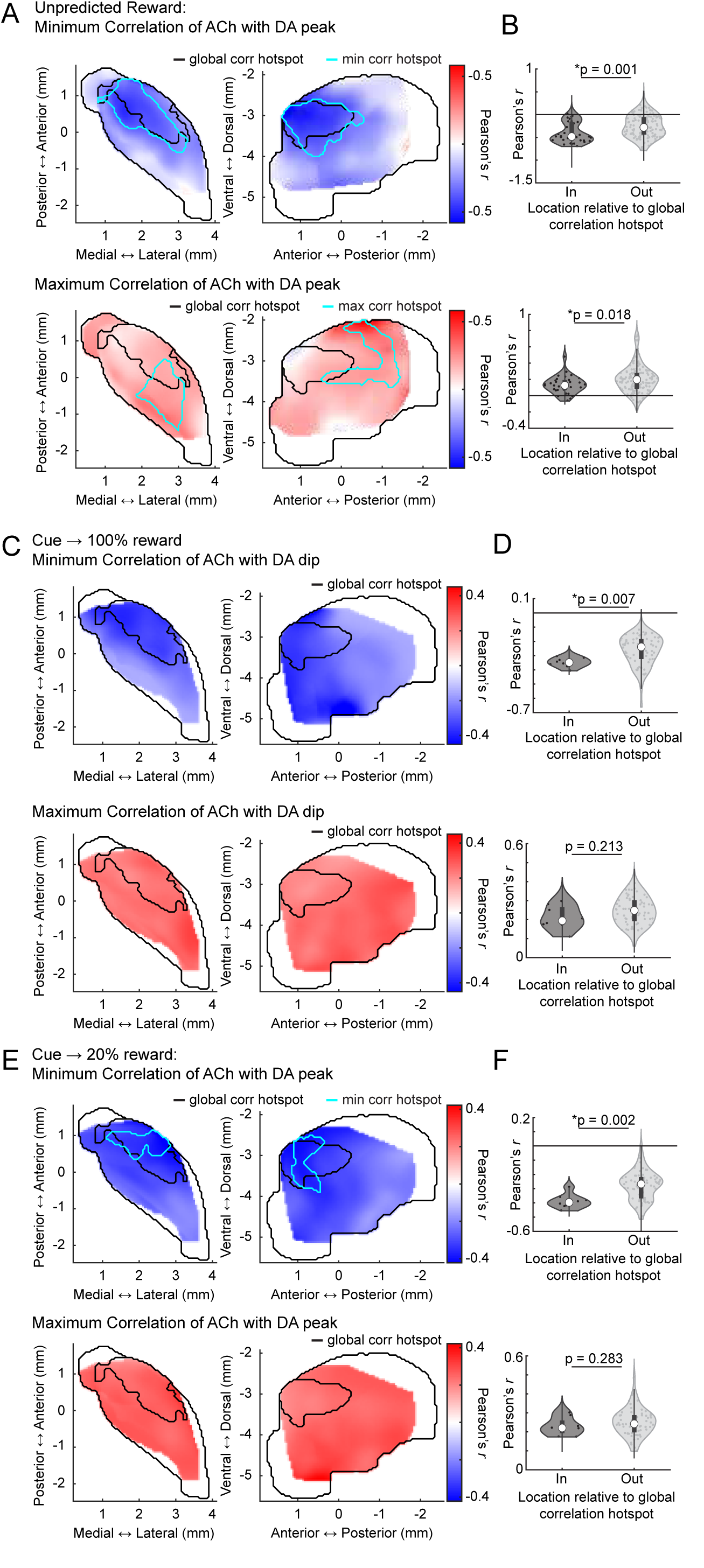
Independent spatial distributions of positive and negative DA-ACh correlations for rewards and conditioned cues. **a,** Smoothed maps (axial, left; sagittal, right) of minimum (top) and maximum (bottom) DA-ACh cross-correlation coefficients (within 1s window) across striatum sites for unpredicted reward deliveries. Black contours delineate the global cross-correlation hotspot (Fig.1f) and cyan contours delineate the significant hotspot for cross-correlation coefficient minima and maxima for unpredicted rewards. **b,** Violin plots of correlation coefficient minima (top) and maxima (bottom) for unpredicted reward deliveries for sites inside and outside the global cross-correlation hotspot. Each dot is one site; white circles, median; thick bar, interquartile range; thin lines, 1.5x interquartile range (*p<0.05, Wilcoxon rank-sum test). **c-d,** Same as a-b for conditioned cue periods. No significant hotspots were identified for either correlation minima or maxima, though a dominant correlation (negative) hotspot was present (Fig. 4b). **e-f,** Same as a-b for extinction cue periods. No significant hotspot was present for correlation maxima. Note that across conditions, correlation minima, but not necessarily maxima, were stronger inside the global cross-correlation hotspot.

**Extended Data Figure 6:**
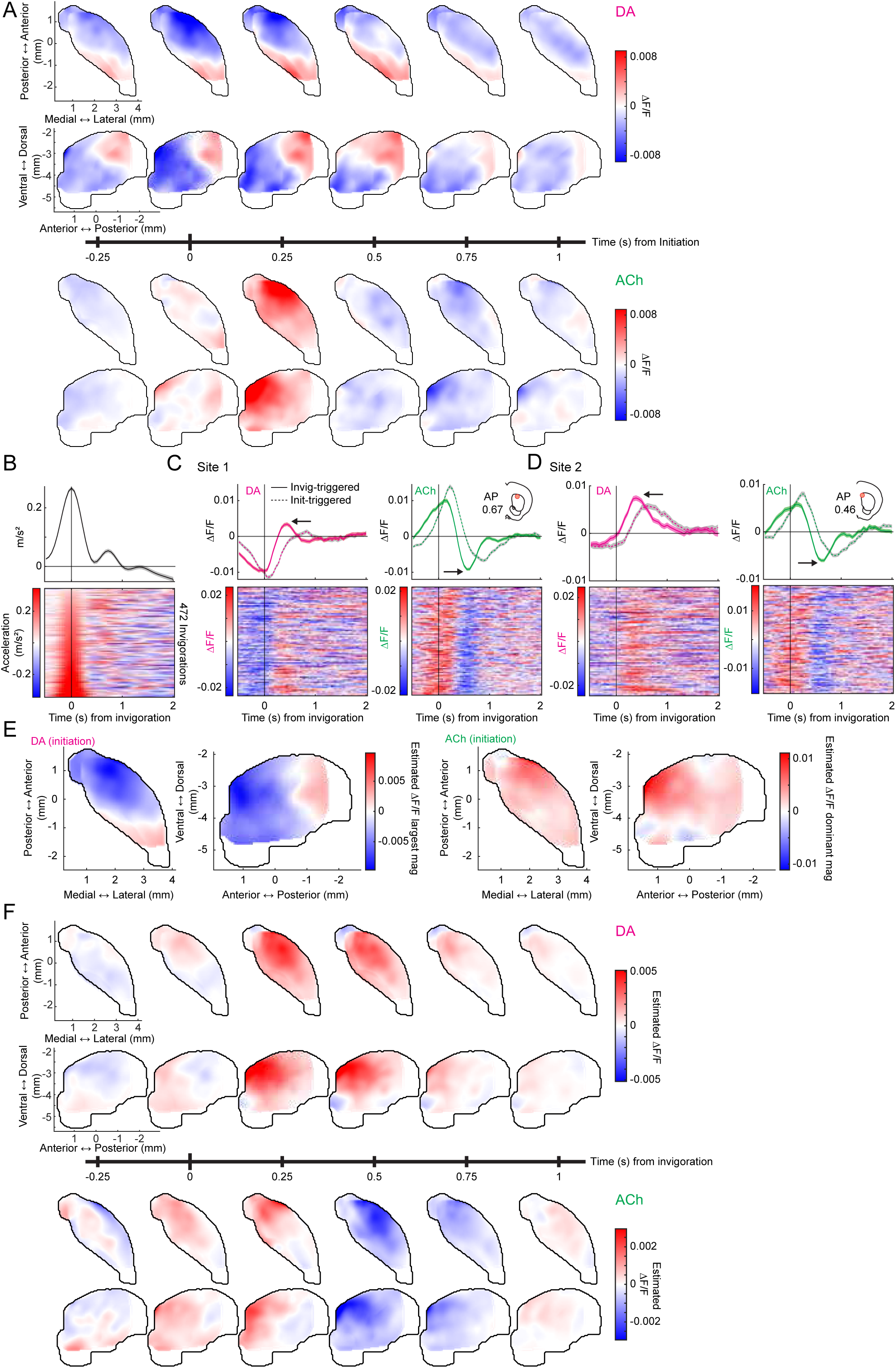
Spatiotemporal features of initiation and invigoration related DA and ACh dynamics. **a,** Smoothed axial and sagittal maps of DA (top two rows) and ACh (bottom two rows) ΔF/F at 0.25s intervals from -0.25 to 1s from locomotion initiations. **b,** Mean (top) and trial-by-trial (bottom) treadmill acceleration aligned to locomotion invigorations for a single mouse, sorted by maximum acceleration. **c,** Mean (top) and trial-by-trial (bottom) DA (left) and ACh (right) ΔF/F for a single example site (location in inset), aligned to invigoration timepoints in b. Dashed lines indicate the mean ΔF/F aligned to the initiation timepoints of the same events. Note that DA peaks and ACh troughs (arrows) are stronger for invigoration alignment, while DA dips and ACh peaks are stronger for initiation alignment. **d,** Same as c for another example site in the same mouse. **e**, Dominant initiation-triggered ΔF/F magnitude estimated from the GLM. Note that the GLM successfully recapitulates the actual dominant initiation-triggered ΔF/F magnitude (Fig. 5e). **f,** Same as a but for invigoration aligned ΔF/F estimated from the GLM.

**Extended Data Figure 7:**
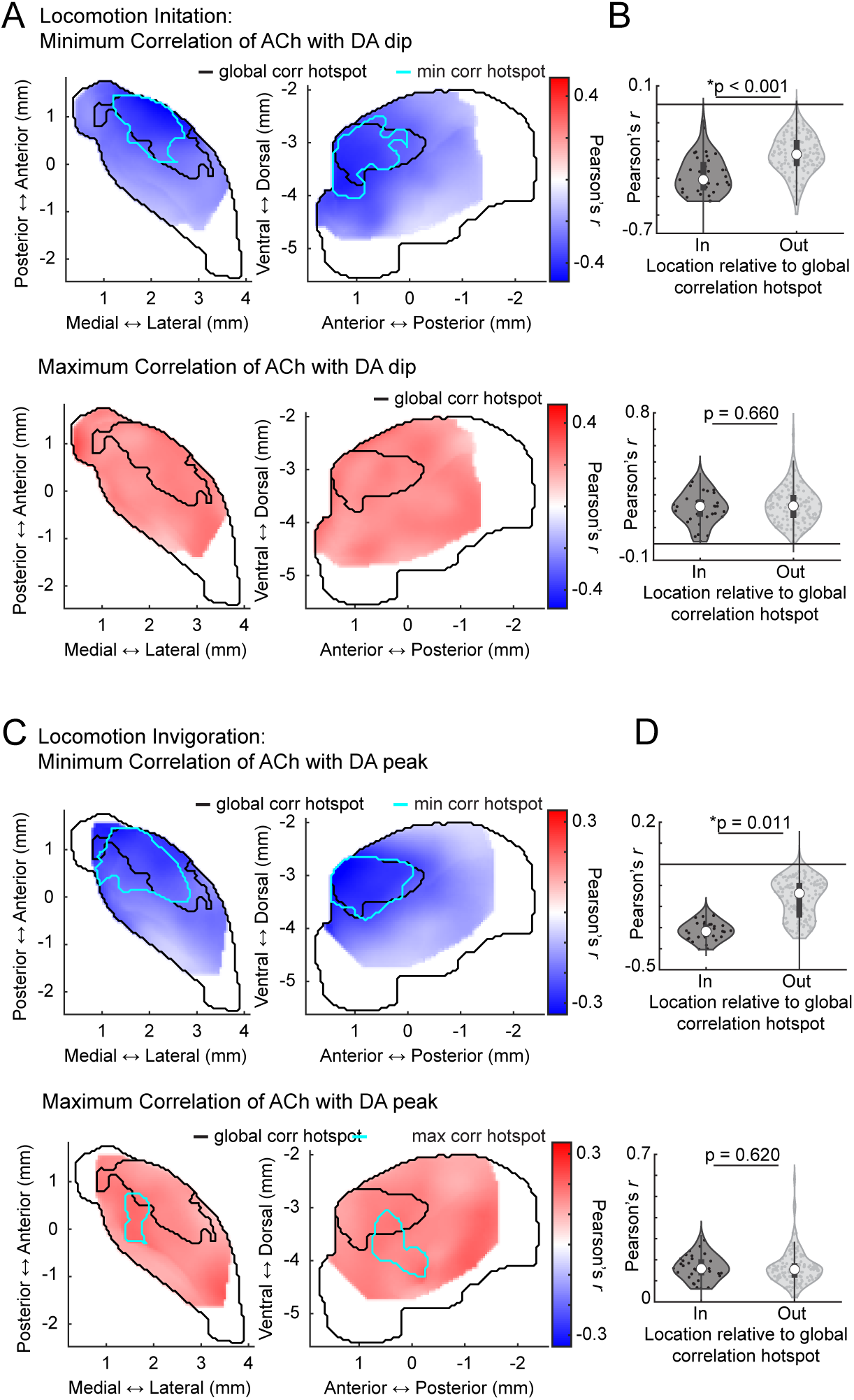
Independent spatial distributions of positive and negative DA-ACh correlations for locomotion initiations and invigorations. **a,** Smoothed maps (axial, left; sagittal, right) of minimum (top) and maximum (bottom) DA-ACh cross-correlation coefficients (within 1s window) across striatum sites for locomotion initiations. Black contours delineate the global cross-correlation hotspot (Fig.1f) and cyan contours delineate the significant hotspot for cross-correlation coefficient minima and maxima for locomotion initiations. No significant hotspot was identified for correlation maxima. **b,** Violin plots of correlation coefficient minima (top) and maxima (bottom) for locomotion initiations for sites inside and outside the global cross-correlation hotspot. Each dot is one site; white circles, median; thick bar, interquartile range; thin lines, 1.5x interquartile range (*p<0.05, Wilcoxon rank-sum test). **c-d,** Same as a-b for locomotion invigorations. Note that for both initiations and invigorations, correlation minima, but not maxima, were stronger inside the global cross-correlation hotspot.

**Extended Data Figure 8:**
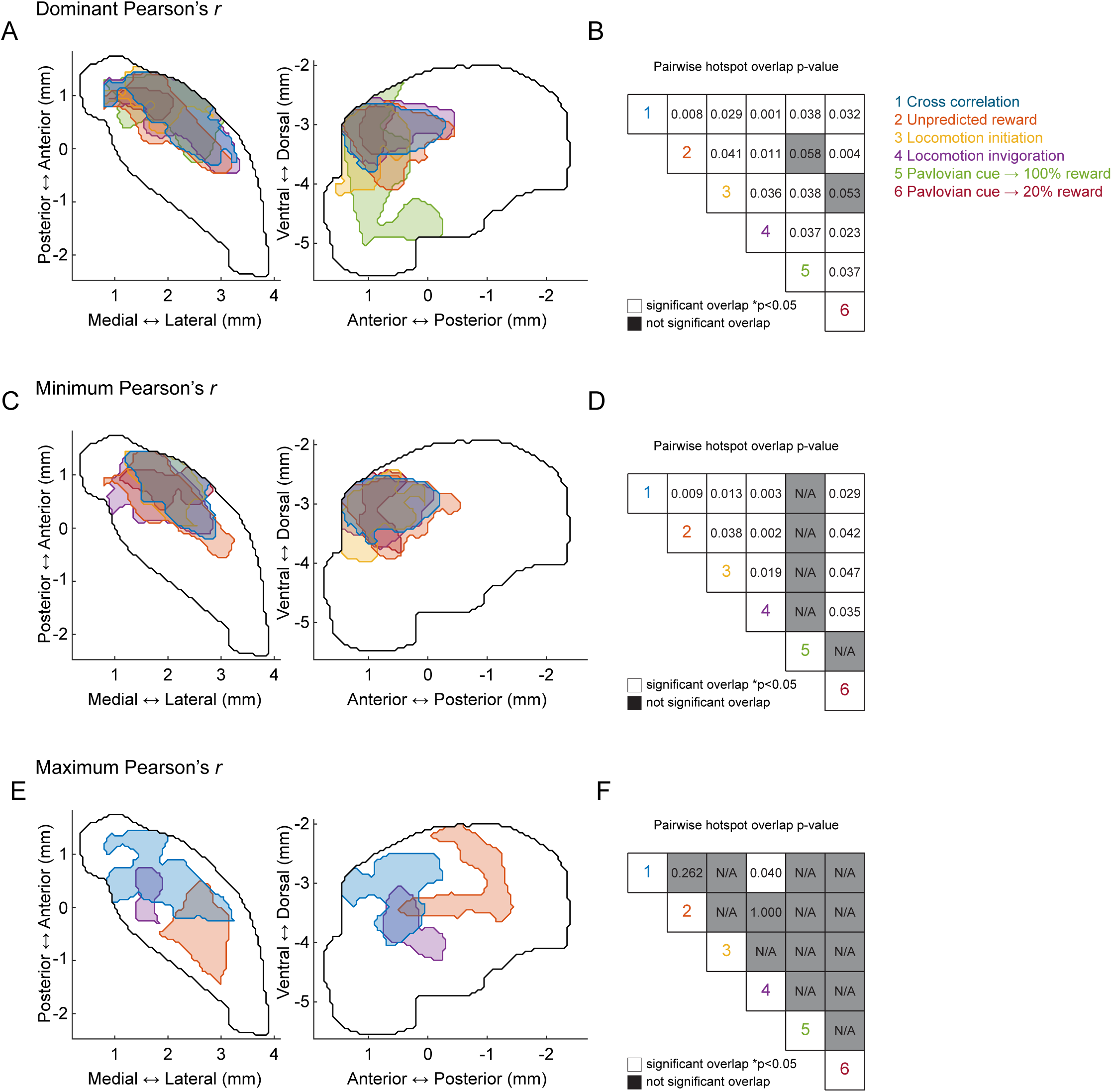
aDLS-concentrated DA-ACh dominant and negative, but not positive, correlation hotspots are consistent. **a,** Maps (axial, left; sagittal, right) of dominant correlation hotspots. Each contour represents the hotspot defined by the dominant correlation analysis in a different experiment, according to the color key in b: 1) blue, cross correlation; 2) orange, unpredicted reward; 3) yellow, locomotion initiation; 4) purple, locomotion invigoration; 5) green, pavlovian cues; 6) red, extinguished pavlovian cues.**b,** Pairwise p-values of overlap between hotspots. White squares indicate significant overlap evaluated at p<0.05. Gray squares indicate non-significant overlap or no identified hotspot (N/A). Note that all hotspots significantly overlap with the global dominant cross-correlation hotspot (top row). **c-d,** same as a-b, but for minimum correlation. Note the significant overlap between all identified minimum correlation hotspots (no minimum correlation hotspot was identified for Pavlovian cues). **e-f**, same as a-b, but for maximum correlation. Note that maximum correlation hotspots were only identified for cross-correlation (blue), unpredicted reward (orange), and locomotion invigoration (purple). The only significant overlap is between the maps for maximum cross-correlation and maximum DA-ACh correlation coefficient in locomotion invigorations.

**Extended Data Figure 9:**
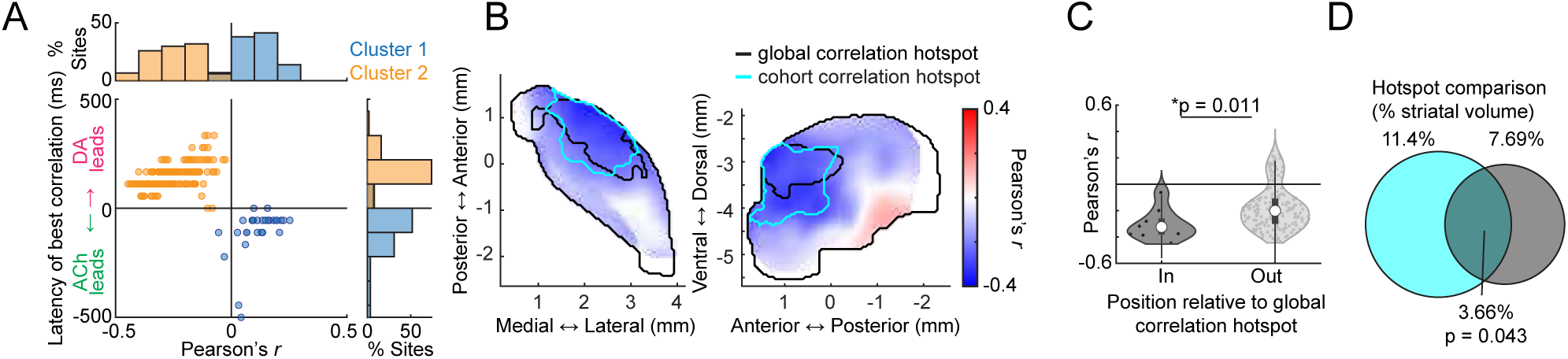
Replication of cross-correlation hotspot dynamics in the DA stimulation cohort. **a,** Dominant DA-ACh correlation coefficient vs latency for each site (dot) across all mice from the DA stimulation cohort (n = 6 mice, 180 sites) for spontaneous release during non-stimulation periods. Clusters identified by k-means. Histograms show the percent of total sites in each cluster binned across correlation coefficients (top) and latencies (right). **b,** Smoothed maps (axial, left; sagittal, right) of dominant DA-ACh correlation coefficients (max or min). Black contours delineate the global cross-correlation hotspot, (original cohort, Fig. 1f) and cyan contours delineate the significant dominant cross-correlation hotspot identified in the DA stimulation cohort. **c,** Violin plots showing dominant DA-ACh cross-correlation coefficients for each site inside or outside the original aDLS global correlation hotspot for the DA stimulation cohort. Each dot is one site; white circles, median; thick bar, interquartile range; thin lines, 1.5x interquartile range. P-values, Wilcoxon rank-sum test; *p<0.05. **d,** Venn diagrams showing the spatial overlap between the global cross-correlation hotspot (black contour, Fig. 1f) and the correlation hotspot in the DA stimulation cohort (cyan contours in b). Percentages indicate the fraction of the total single-hemisphere striatum volume occupied by individual hotspots (cyan left, black right) and for the intersection of the two hotspots (middle, volume overlap test, see Methods).

**Extended Data Figure 10:**
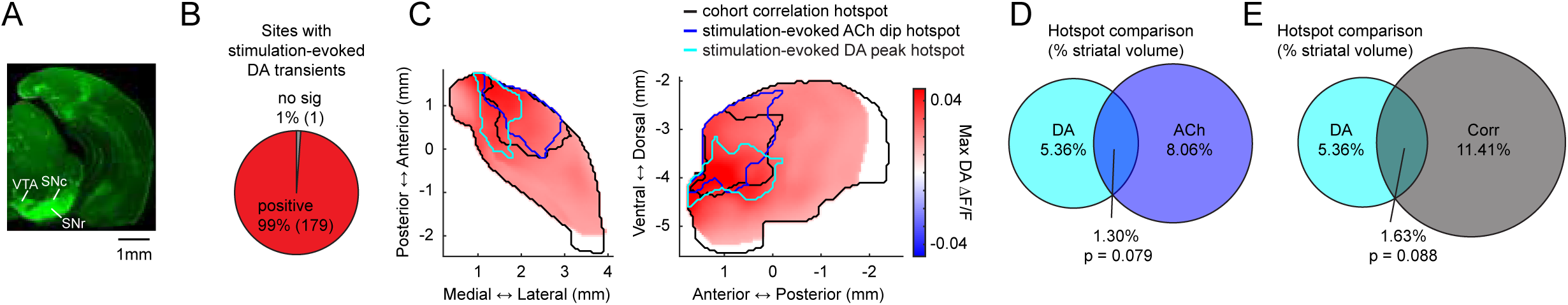
Spatial biases in stimulation-evoked DA release magnitudes do not account for hotspot localized ACh effects. **a,** Representative coronal slice showing immunolabeling confirmation of expression of ChR2 in midbrain. VTA, ventral tegmental area; SNc, substantia nigra pars compacta; SNr, substantia nigra pars reticulata. **b,** Fraction of sites with significant stimulation-evoked DA release. **c,** Smoothed maps (axial, left; sagittal, right) of mean stimulation-evoked DA ΔF/F peak magnitudes. Contours indicate the spontaneous anti-correlation hotspot (black), the hotspot of ACh dips evoked by DA stimulation (blue), and the hotspot of stimulation-evoked DA peaks (cyan). **d**, Venn diagram showing the spatial overlap between the DA stimulation-evoked ACh dip hotspot (blue contour in e) and the stimulation-evoked DA peak hotspot (cyan contour in e). Percentages indicate the fraction of the total single-hemisphere striatum volume occupied by individual hotspots and for the intersection of the two hotspots (middle, volume overlap test, see Methods). **e,** Same as g but comparing the stimulation-evoked DA peak hotspot (cyan) to the spontaneous anti-correlation hotspot during non-stimulation periods (black). Note that the stimulation-evoked ACh dip hotspot (blue) and correlation hotspot (black) overlap more with each other (Fig. 7g) than with the stimulation-evoked DA peak hotspot (cyan), with which neither significantly overlaps.

